# Oleic acid suppresses NLRP3 inflammasome activation to prevent dermal macrophage death and limit imiquimod-induced psoriasis

**DOI:** 10.64898/2026.10.27.760067

**Authors:** Beatriz Burger, Roberta Nicolli Sagiorato, Isabela Buquio Baccarin, Vitor Rosetto Muñoz, Gabriella Cerna, Ilya Andreev, Nathália Araújo, Hui Sophie Shu, Jia Zhao, Brandon Sullivan, Yisha Liang, Rodrigo Angelo Martinati Bistaffa, Vitória C. Gaspar, Esther Midori Yabuuti, Natalie Bitencourt Ramos, Laís Passariello Pral, Charles Evavold, José Carlos Alves Filho, Patrick Varga-Weisz, Carlos Henrique Cardoso Serezani, Marco Aurélio Ramirez Vinolo, Hernandez Moura Silva, Hosana Gomes Rodrigues

**Author notes:** Corresponding author: Hosana Gomes Rodrigues. These authors contributed equally to this work (Hernandez Moura Silva, Hosana Gomes Rodrigues).

## Abstract

Psoriasis is associated with altered lipid metabolism, including reduced expression of fatty acid metabolism genes and decreased circulating oleic acid. Here we report that oral oleic acid supplementation normalized serum oleic acid levels and protected from skin thickening and epidermal hyperplasia in a imiquimod mice model of psoriasis. These effects were associated with preserved skin-resident macrophage numbers and reduced expression of key psoriasis-associated inflammatory markers. Mechanistically, imiquimod induces dermal macrophage death via NLRP3/caspase-1 inflammasome activation, while adoptive transfer of dermal macrophages attenuates psoriasiform inflammation. Oleic acid rescued macrophages from imiquimod and nigericin-induced death *in vitro* by suppressing NF-κB and NLRP3/caspase-1 signaling, lowering IL-1β and reactive oxygen species production, and restoring mitochondrial homeostasis, independently of TLR7. Together, our results position dermal macrophage survival as a previously unrecognized checkpoint in psoriasiform inflammation, and suggest that restoring lipid metabolic fitness, rather than solely targeting downstream inflammatory mediators, may offer an additional mechanism-based strategy to limit tissue-damaging inflammation.

## INTRODUCTION

Psoriasis is one of the most prevalent immune-mediated inflammatory skin disorders, affecting approximately 2–3% of the global population (1–4). The disease is defined by epidermal hyperplasia, aberrant keratinocyte activation, and persistent immune cell infiltration into the skin, which together sustain a chronic, self-amplifying inflammatory state (5,6). Among the resident immune populations that shape this chronicity, macrophages occupy a central position: they maintain skin homeostasis through phagocytosis, antigen presentation, and cytokine production, and recent work has revealed remarkable functional heterogeneity among skin macrophage populations, alongside their critical contribution to the initiation and maintenance of inflammatory skin disease (7–9).

This protective function is disrupted in psoriatic skin in both mice and humans, where dermal resident macrophages (DMs) undergo metabolic and mitochondrial alterations, including disrupted mitochondrial respiration and excess reactive oxygen species (ROS) generation, that drive NLRP3 inflammasome activation and inflammatory cell death, amplifying innate immune responses and sustaining the IL-23/IL-17 axis that underlies chronic disease (10–15). These findings posit dermal macrophage death as a self-amplifying node in psoriasis pathogenesis, underscoring the need to identify the upstream signals that determine whether these macrophages survive or succumb to this inflammatory environment. Because metabolic and inflammatory processes in immune cells can be influenced by nutrient availability, dietary components represent potential upstream regulators of macrophage responses during skin inflammation. Among these, fatty acids are of particular interest because dietary fatty acid intake can alter the availability of lipid substrates and signaling molecules that regulate cellular metabolism and inflammatory pathways (16). They shape immune cell proliferation, differentiation, survival, and inflammatory signaling by modulating cellular metabolism, membrane composition, and intracellular signaling pathways (17–21). Fatty acids are classified by the number of double bonds in their carbon chain as saturated, monounsaturated, or polyunsaturated and their immunomodulatory effects vary accordingly (20,22). Oleic acid (OA) is an mono-unsaturated omega-9 fatty acid that is both endogenously synthesized and obtained from the diet, most notably from olive oil, a hallmark of the Mediterranean diet. OA attracted particular attention for its anti-inflammatory properties and its association with reduced risk of metabolic and chronic inflammatory disease (23,24). Supporting a broader role for diet in shaping disease outcomes, experimental and clinical studies indicate that dietary fatty acid intake can modulate the susceptibility and severity of inflammatory conditions (16,25).

We previously showed that oral supplementation with OA improves wound healing in diabetic rats, pointing to immunomodulatory and tissue-protective properties beyond its established metabolic effects (26–30). In parallel, dysregulated lipid metabolism has emerged as a feature of psoriasis, contributing to immune cell activation and inflammatory cytokine production (16,31). However, whether modulating OA levels, through either endogenous synthesis or dietary supplementation, could mitigate psoriasis pathology had not been tested.

Here, we show that dietary OA supplementation attenuates skin inflammation and disease severity in the imiquimod-induced mouse model of psoriasis by protecting dermal macrophages from NLRP3-dependent inflammatory cell death, establishing a protective immunometabolic role for dietary oleic acid in psoriasis.

## RESULTS

### Reduced fatty acid biosynthesis and circulating oleic acid levels in human and murine psoriasis

To identify canonical inflammatory pathways mediating psoriasis, we defined the transcriptional signature of human psoriatic skin and asked whether it was recapitulated in the imiquimod-induced mouse model of psoriasiform inflammation. Analysis of three independent public datasets of lesional versus healthy skin (32–34), comprising a total of 187 and 160 samples, respectively, identified 251 genes consistently upregulated across psoriatic lesions **(Supplementary Figure 1A)**, forming a signature enriched for inflammatory mediators, including members of the S100A family, interleukins, and chemokines **(Supplementary Figure 1B-C)**, consistent with previous transcriptomic studies of psoriasis (35,36).

To determine whether this signature was conserved in experimental psoriasis, we characterized the imiquimod-induced mouse model **(Supplementary Figure 1D)** (37). Four consecutive days of imiquimod treatment induced hallmark features of psoriasis-like skin inflammation, including epidermal hyperplasia, increased skin thickness, erythema, and scaling **(Supplementary Figure 1E-H)**, alongside systemic manifestations such as body weight loss and increased spleen and lymph node weight **(Supplementary Figure 1I-J)** (38,39). Temporal analysis of psoriasis-associated gene expression identified a peak inflammatory response after 3 days of imiquimod treatment **(Supplementary Figure 1K)**, the time point selected for RNA sequencing. Transcriptomic profiling of whole skin at this time point revealed a robust inflammatory response, including the upregulation of genes associated with psoriasis and innate immune activation **(Supplementary Figure 1L-M)**. Together, these findings show that the imiquimod-induced mouse model recapitulates both the phenotypic and core inflammatory transcriptional features of human psoriasis.

Given that skin is an important lipid reservoir and fatty acids are key regulators of cutaneous immune responses, we next asked whether lipid metabolism genes were similarly dysregulated in the three human datasets **(Figure 1A)**. Functional enrichment analysis show that genes involved with lipid biosynthesis, fatty acid metabolism and lipid metabolism where downregulated across the three psoriasis datasets **(Figure 1B)**, indicating a marked suppression of lipid metabolic programs in psoriatic skin. Notably, expression of stearoyl-CoA desaturase 5 (SCD5), negatively correlated with the psoriasis markers S100A8 and S100A9 **(Figure 1C)**. Imiquimod-treated mice recapitulated this lipid phenotype, showing reduced expression of the lipid biosynthesis genes *Scd3*, *Elovl6*, *Fads2*, and *Scd4* **(Figure 1D-E)**. As SCD is the endoplasmic reticulum membrane enzyme that catalyzes desaturation of stearic acid to generate oleic acid **(Figure 1F),** we reasoned that this suppression would lower circulating levels of oleic acid, and indeed we observed a 40.5% decrease in serum oleic acid and a 35.5% decrease in serum linoleic acid in imiquimod-treated mice **(Figure 1G).** Daily food intake did not differ between control and imiquimod-treated mice **(Figure 1H)**, indicating that these reductions were not attributable to decreased food consumption. Although we observed significant reductions in oleic and linoleic acid levels, no changes were detected in other fatty acid classes, including omega-3 fatty acids **(Figure 1G),** indicating that this effect may be selective rather than a nonspecific alteration in fatty acid level.

**Figure 1.**
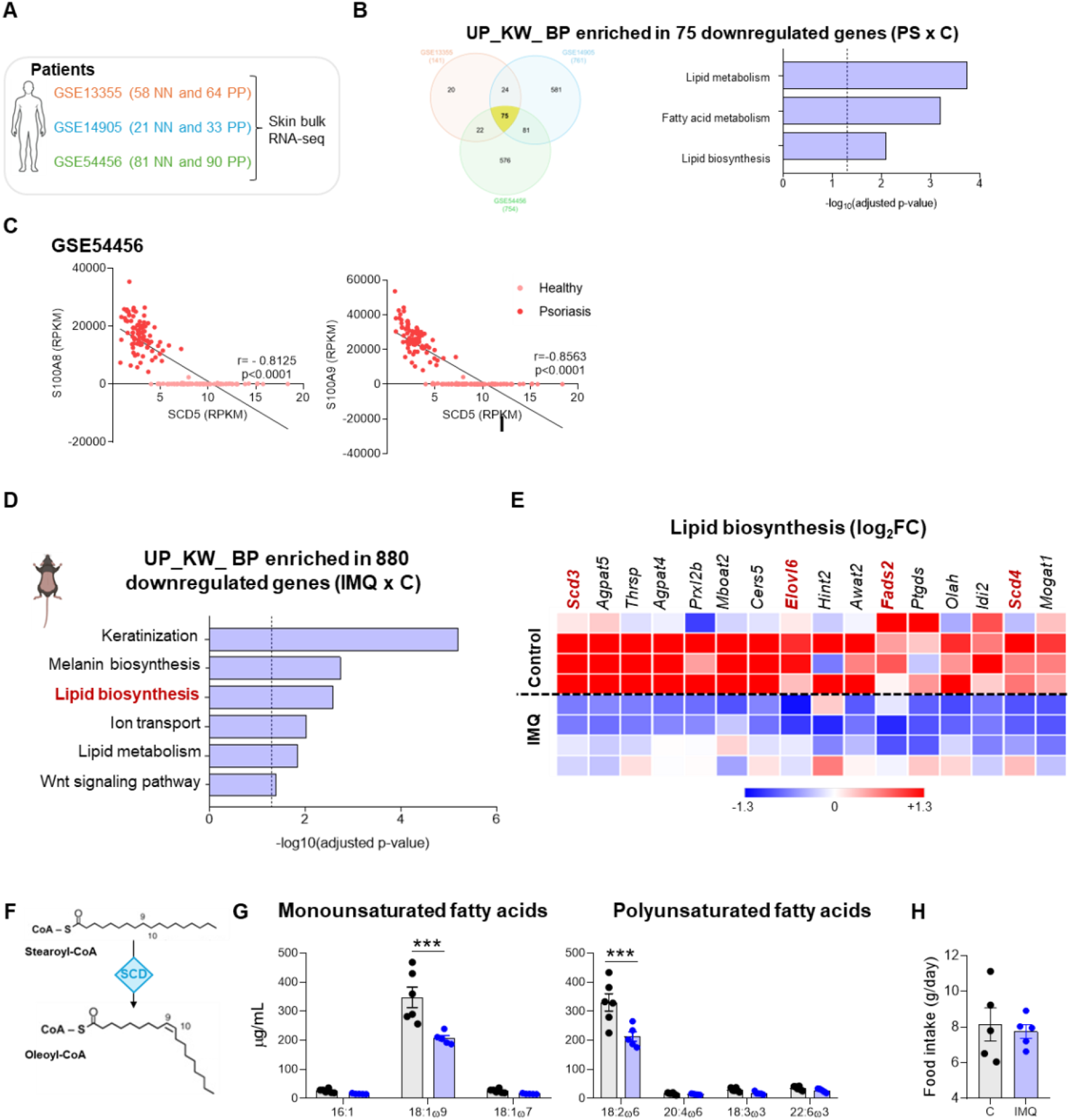
Human and murine psoriasis exhibit impaired fatty acid metabolism. **(A)** Human datasets samples. **(B)** Venn diagram showing the overlap of downregulated genes identified across three human psoriasis datasets, considering log_2_(fold change) ≥ 1.3 or ≤−1.3 and adjusted p-value<0.05 and gene ontology (GO) of Biological Processes enriched in 75 commonly downregulated genes in psoriatic versus healthy skin across the three datasets. **(C)** Correlation analysis between *SCD5* expression and psoriasis-associated markers (*S100A8* and *S100A9*), measured as Reads Per Kilobase Million (RPKM), in 81 healthy human skin samples and 90 psoriatic skin samples from the GSE54456 dataset, assessed using Spearman’s correlation test. **(D)** Top downregulated Biological Process identified in the skin following imiquimod treatment. **(E)** Heatmap of genes associated with lipid biosynthesis pathways. Each row represents an individual mouse and each column represents a gene. **(F)** Schematic illustration of SCD-mediated conversion of stearoyl-CoA into oleoyl-CoA. **(G)** Serum levels of monounsaturated and polyunsaturated fatty acids in mice. **(H)** Daily food intake in control and imiquimod-treated mice (g/mouse/day). Values are expressed as mean ± SEM. (***) p≤0.001 by two-way ANOVA followed by Bonferroni’s post hoc test (G) or unpaired Student’s t-test (H)

Together, these findings reveal a conserved suppression of lipid metabolic programs in human psoriatic skin and in the imiquimod-induced mouse model, marked by reduced expression of lipid biosynthesis and desaturation enzymes and decreased circulating oleic acid levels. These results indicate that psoriasis is associated with impaired endogenous fatty acid metabolism in the skin, and support the hypothesis that modulating oleic acid levels may ameliorate disease pathophysiology.

### Oral oleic acid supplementation attenuates psoriasiform inflammation and preserves resident tissue macrophages in the skin

To determine whether oral oleic acid supplementation restores circulating oleic acid levels, we supplemented imiquimod-treated mice with pure oleic acid with a dose corresponding to approximately 0.14 g/kg/day in humans based on body surface area scaling, or ∼10 g/day for a 70-kg adult **(Figure 2A)** (40), within the range of habitual dietary oleic acid intake reported in human populations (41). Oral supplementation normalized serum oleic acid levels in mice with imiquimod-induced psoriasis **(Figure 2B)**, Compared with imiquimod-treated mice, mice receiving oral oleic acid during imiquimod treatment exhibited reduced back skin thickness and, macroscopically, showed less severe inflammation **(Figure 2C-D),** as well as decreased epidermal thickness by H&E staining **(Figure 2E-F).** Body weight remained comparable among groups throughout the experimental period, with no significant effect of oleic acid supplementation **(Supplementary Figure 2A).** The protective effect of oral oleic acid supplementation was also observed in female mice, in which supplementation attenuated the increase in skin thickness following imiquimod treatment without affecting body weight **(Supplementary Figure 2B-F).**

**Figure 2.**
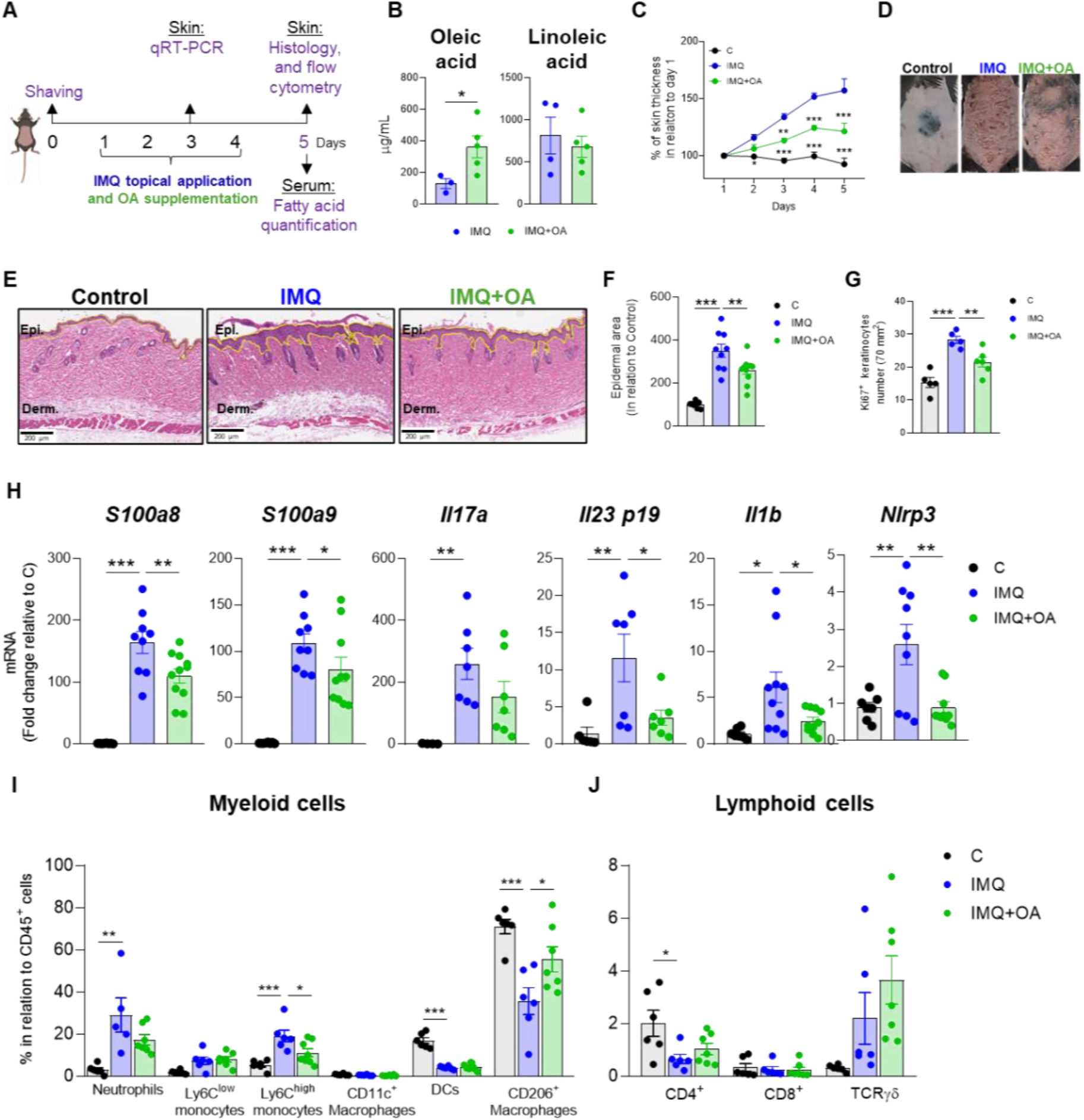
Dietary Oleic Acid attenuate imiquimod-induced psoriasis in mice and preserved the resident dermal macrophages. **(A)** Experimental design: dorsal skin was shaved one day before psoriasis induction, followed by daily topical application of 60 mg of 5% imiquimod cream from day 1 to day 4, concomitant with oleic acid (OA) supplementation. Skin thickness was monitored daily, and skin samples were collected on day 3 for qRT-PCR or on day 5 for histological and flow cytometry analyses. **(B)** Serum concentrations of oleic and linoleic acid. **(C)** Percentage of skin thickness relative to day 1. **(D)** Representative photographs of mouse dorsal skin. **(E)** Representative histological images of dorsal skin collected on day 5. Scale bar: 200 µm. **(F)** Quantification of epidermal area relative to the control group. **(G)** Quantification of Ki67⁺ keratinocytes in the epidermis. **(H)** Relative gene expression of *S100a8*, *S100a9*, *Il17a*, *Il23p19*, *Il1β* and *Nlrp3* in skin samples collected on day 3. **(I, J)** Percentage of immune cells in relation to CD45^+^ cells after 5 days. Values are expressed as mean ± SEM. (*) p≤0.05; (**) p≤0.01; (***) p≤0.001 by unpaired Student’s t-test (B) or two-way ANOVA (C, I, J) or by one-way ANOVA followed by Dunnett’s post hoc test (F, G, H).

Consistent with reduced epidermal hyperproliferation, staining for the proliferation marker Ki67 and the basal keratinocyte marker K14 showed that imiquimod treatment markedly increased Ki67+ cells within the basal and, to a lesser extent, suprabasal keratinocyte layers, whereas oleic acid supplementation reduced the number of Ki67^+^ basal cells relative to imiquimod alone **(Figure 2G and Supplementary Figure 2G).**

To determine whether this phenotypic improvement reflected a corresponding change in the molecular drivers of psoriatic inflammation, we profiled expression of genes classically associated with disease severity. Oleic acid supplementation inhibited the upregulation of key genes in the skin of imiquimod-treated mice **(Figure 2H).**

To further characterize the effects of oleic acid supplementation on the skin immune compartment, we analysed the composition of dermal immune cell populations. Neutrophils and Ly6C^high^ monocytes were strongly increased following imiquimod treatment **(Figure 2I)**, indicating enhanced recruitment of circulating myeloid cells to the inflamed tissue. In contrast, CD11c⁺ macrophages were not substantially altered by imiquimod, while dendritic cells and CD206⁺ macrophages were markedly reduced **(Figure 2I).** Notably, OA supplementation partially protected the skin from these changes, suppressing Ly6C^high^ monocytes accumulation and preserving the CD206⁺ macrophage population. **(Figure 2I).** Within the lymphoid compartment, imiquimod inhibited the accumulation of CD4⁺ T cells, whereas CD8 and TCRγδ⁺ T cells were not affected **(Figure 2J).**

Notably, CD206^+^ dermal macrophages (DMs) represent a population of resident tissue macrophages associated with skin homeostasis and repair (42,43). Resident tissue macrophages are essential regulators of organ homeostasis, coordinating repair responses and supporting specialized tissue functions (7,9). These cells are maintained through a dynamic balance between local self-renewal and monocyte recruitment, and disruption of their abundance or functional capacity has been associated with several chronic inflammatory and age-related diseases, including atherosclerosis, cancer, and neurodegeneration (44). Considering the marked reduction in resident DMs during psoriasiform inflammation and their preservation in the skin of mice receiving oral oleic acid supplementation, we next investigated the dynamics of macrophage replenishment during disease progression and the effects of oleic acid supplementation and used an established procedure in our lab to functionally fate-map DMs and track these cells throughout disease progression **(Figure 3A).** Intravenous injection of dextran-rhodamine prior to psoriasis induction selectively and stably labels resident DMs already present in the skin, while sparing circulating monocytes that have not yet been recruited to the tissue. Because newly recruited monocyte-derived macrophages are unlabelled, retention of dextran-rhodamine signal over time indicates persistence of the original resident population, whereas loss of labelling reflects their replacement by monocyte-derived cells. Using this approach, we found that CD206^+^ macrophages are the main population to express dextran-rhodamine **(Figure 3B-C)** and, the majority of DMs was replaced by unlabelled, monocyte-derived cells in the imiquimod group **(Figure 3D-E)**. In contrast, oral oleic acid supplementation prevented the loss of dextran-rhodamine^+^ DMs, indicating that oleic acid preserves the pre-existing resident macrophage compartment during psoriasiform inflammation. These findings raised the question of whether the accumulation of newly recruited macrophages during the inflammatory response resulted from increased replenishment of the tissue by circulating precursors.

**Figure 3.**
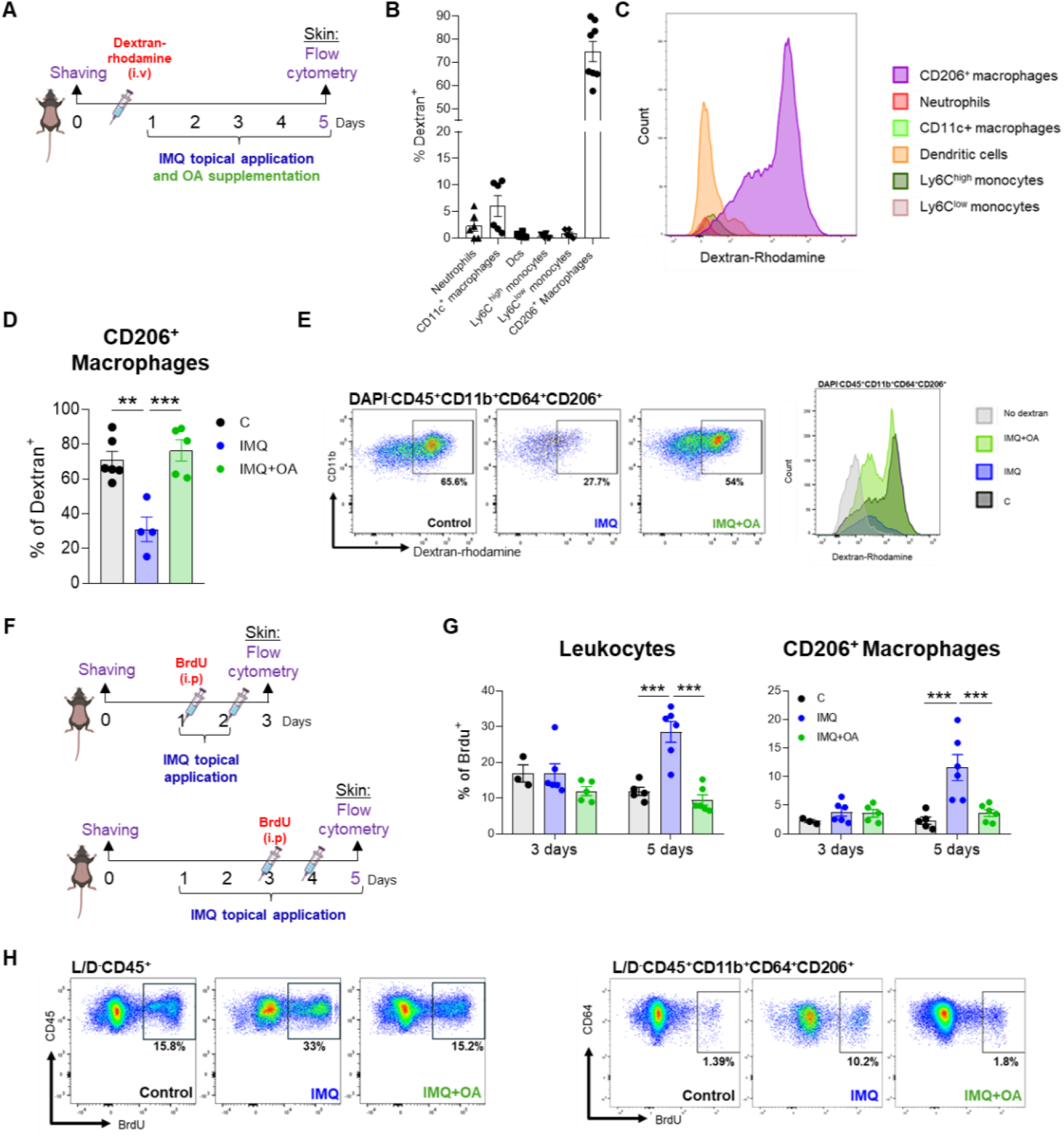
Oleic acid supplementation preserves resident dermal macrophages. **(A)** For macrophages resilience analysis, Dextran-Rhodamine was injected retro-orbitally between days 0 and 1, and skin samples were collected on day 5 for flow cytometry. **(B)** CD206⁺ macrophages (dashed line) exhibit high endocytic activity *in vivo*, as they represent the predominant dextran⁺ cell population. **(C)** Histograms display the fluorescent intensity for rhodamine of skin populations. **(D)** Quantification of Dextran^+^ resident macrophages after 5 days. **(E)** Representative flow cytometry plots of Dextran⁺ resident macrophages. **(F)** Experimental design: dorsal skin was shaved one day before psoriasis induction, followed by daily topical application of 60 mg of 5% imiquimod cream from day 1 to day 2 or 4. For proliferation analysis, Brdu (40 mg/Kg; 100 µL) was administered intraperitoneally every 12 hours on days 1 and 2 or on days 3 and 4, and skin samples were collected on day 3 or 5 for flow cytometry. **(G)** Frequency of BrdU⁺ leukocytes and CD206⁺ macrophages. **(H)** Representative flow cytometry plots showing BrdU⁺ leukocytes and BrdU⁺ CD206⁺ macrophages. Values are expressed as mean ± SEM. (*) p≤0.05; (**) p≤0.01; (***) p≤0.001 by by one-way ANOVA followed by Dunnett’s post hoc test (D) or two-way ANOVA followed by Bonferroni’s post hoc test (G).

To further investigate the cellular dynamics underlying this replenishment, we next used BrdU incorporation to identify cells undergoing DNA synthesis during defined phases of disease progression **(Figure 3F).** Because circulating monocytes can proliferate before their recruitment into inflamed tissues, BrdU labeling provides a means to detect proliferating precursors that subsequently contribute to the macrophage compartment. Mice received intraperitoneal injections of BrdU every 12 hours for two consecutive days during either the early phase (days 1–2) or late phase (days 3–4) of psoriasis induction, and skin samples were collected on days 3 or 5, respectively **(Figure 3F).** Imiquimod treatment increased the proportion of BrdU^+^ leukocytes during the late phase of disease, with macrophages representing the predominant BrdU^+^ population, whereas oleic acid supplementation maintained BrdU^+^ leukocyte and macrophage frequencies at levels comparable to those observed in control mice **(Figure 3G-H).** Together with the dextran-rhodamine fate-mapping results, these findings support a model in which the increase in macrophages during the late inflammatory phase is associated with replenishment of the DM compartment by newly recruited monocyte-derived cells rather than *in situ* proliferation of resident macrophages.

To determine whether restoring dermal macrophages could influence the development of psoriasiform inflammation, we performed adoptive transfer of dermal macrophages intradermally during imiquimod-induced psoriasiform inflammation **(Figure 4A).** Macrophage transfer significantly reduced skin thickness and epidermal area **(Figure 4B-E)** and attenuated the imiquimod-induced increase in *S100a8* expression in the skin after 3 days of treatment **(Figure 4F).** At the same time, we observed increased *Arg-1* expression in the skin, indicating the restoration of resident macrophages following injections **(Figure 4F).** Collectively, these observations suggest that the reduction of resident dermal macrophages contributes to psoriasis development, while maintenance of this macrophage population may attenuate skin inflammation and disease severity.

**Figure 4.**
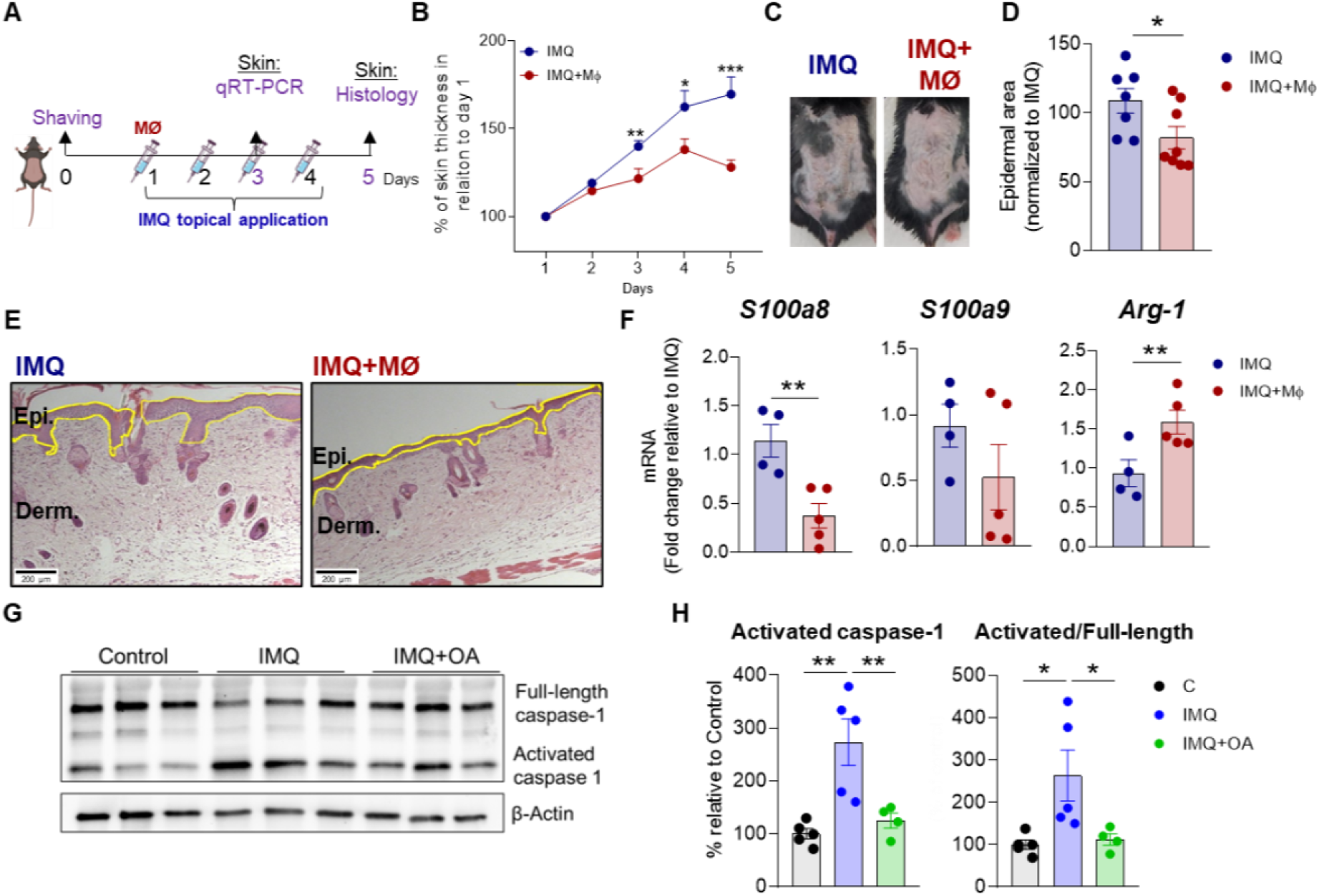
Resident dermal macrophage loss promotes psoriasiform inflammation, whereas macrophage restoration limits disease and oral oleic acid supplementation reduces caspase-1 activation. **(A)** Experimental design for imiquimod-induced psoriasiform inflammation: dorsal skin was shaved one day before disease induction, and mice received daily topical applications of 60 mg of 5% imiquimod cream from day 1 to day 4, concomitantly with subcutaneous macrophage injection. Skin thickness was monitored daily, and skin samples were collected on day 5 for histological analysis or on day 3 for rtPCR. **(B)** Percentage of skin thickness relative to day 1. **(C)** Representative photographs of mouse dorsal skin. **(D)** Quantification of epidermal area relative to the control group. **(E)** Representative histological images of dorsal skin collected on day 5. Scale bar: 200 µm. (F) *S100a8, S100a9* and *Arg-1* mRNA expression in skin samples collected on day 3. **(G)** Representative Western blot analysis of Caspase-1**. (H)** Quantification of activated caspase-1 and cleaved Caspase-1/full-length Caspase-1 ratio. Values are expressed as mean ± SEM. (*) p≤0.05; (**) p≤0.01; (***) p≤0.001 by by two-way ANOVA followed by Bonferroni’s post hoc test (B) or by unpaired Student’s t-test (D, F) or by one-way ANOVA followed by Dunnett’s post hoc test (H).

Imiquimod is known to promote TLR7/8 activation, but it also exerts receptor-independent effects (14). These effects have been associated with increased intracellular calcium (Ca^2+^) levels through the IP3 receptor (45), as well as inhibition of mitochondrial complex I, enhanced ROS production, increased mitochondrial membrane potential, NLRP3 inflammasome assembly, leading to caspase-1 activation, IL-1β maturation, and inflammatory myeloid cell death (14) **(Supplementary Figure 3A).** Given that caspase-1 is a central effector of inflammasome signaling, mediating both cytokine processing and inflammatory cell death, we next assessed caspase-1 activation in the skin. As expected, imiquimod increased the levels of cleaved-to-full-length caspase-1 ratio **(Figure 4G-H)**, supporting the involvement of inflammasome activation in the induction of cell death. The preservation of macrophages in oleic acid-supplemented mice was accompanied by a reduction in the cleaved-to-full-length caspase-1 ratio to levels comparable to those observed in untreated controls **(Figure 4G-H)**, indicating that oleic acid limits inflammasome activation *in vivo*. We therefore hypothesized that oral oleic acid supplementation could protect dermal macrophages from inflammation-induced cell death during imiquimod treatment. As oleic acid supplementation reduced caspase-1 activation in the skin, we next investigated whether caspase-1 activity contributes to imiquimod-induced psoriasiform inflammation. Pharmacological inhibition of caspase-1 with VX765 **(Supplementary Figure 3B)** markedly attenuated imiquimod-induced psoriasiform inflammation, as evidenced by reduced skin thickness, macroscopic disease severity, histopathological changes, and gene expression of hallmarks of psoriasis **(Supplementary Figure 3C-G).** Collectively, these findings show that oral oleic acid supplementation attenuates imiquimod-induced psoriasiform inflammation while preserving resident dermal macrophages. This effect was associated with reduced macrophage loss, decreased accumulation of monocyte-derived inflammatory macrophages during the late inflammatory phase, and reduced caspase-1 activation in the skin. Moreover, adoptive transfer of dermal macrophages attenuated key features of psoriasiform inflammation, supporting a functional contribution of the dermal macrophage compartment to disease progression and suggesting that preservation of these cells may contribute to the protective effects of oleic acid.

### Oleic acid suppresses NLRP3 inflammasome activation inhibiting oxidative stress and macrophage cell death

Having established that caspase-1 activation and the loss of resident dermal macrophages contribute to imiquimod-induced psoriasiform inflammation *in vivo*, we next asked whether imiquimod directly triggers inflammasome-dependent macrophage death, using two complementary *in vitro* models, a dermal macrophage lineage (DMLs) and bone marrow-derived macrophages (BMDMs). Cells were pretreated with the NLRP3 inhibitor, MCC950, or the caspase-1 inhibitor, VX765, prior to and throughout imiquimod stimulation *in vitro* **(Supplementary Figure 4A).** Pharmacological inhibition of either NLRP3 or caspase-1 completely abolished imiquimod-induced cell death, indicating that this process is dependent on NLRP3 inflammasome and caspase-1 activation **(Supplementary Figure 4B-C).** Pre-treatment with oleic acid protected both DMLs **(Figure 5A-C)** and BMDMs **(Supplementary Figure 4D-E)** from imiquimod-induced cell death, establishing a cytoprotective effect *in vitro*. Oleic acid’s protection was also cytokine-selective, suppressing the imiquimod-induced rise in IL-1β in DMLs while leaving TNF-α production unaffected **(Figure 5D).** Oleic acid also reduced IL-1β when present during LPS priming alone, consistent with attenuated TLR4 signaling **(Supplementary Figure 4F).** This priming effect, however, could not fully explain the protection against cell death. To distinguish effects on TLR4-dependent priming from effects during subsequent inflammasome activation, cells were first primed with LPS, washed to remove LPS, and then treated with oleic acid during imiquimod exposure. Under these conditions, oleic acid still reduced cell death **(Supplementary Figure 4G-H)**, indicating that its protective effect is not solely attributable to inhibition of TLR4-dependent priming.

**Figure 5.**
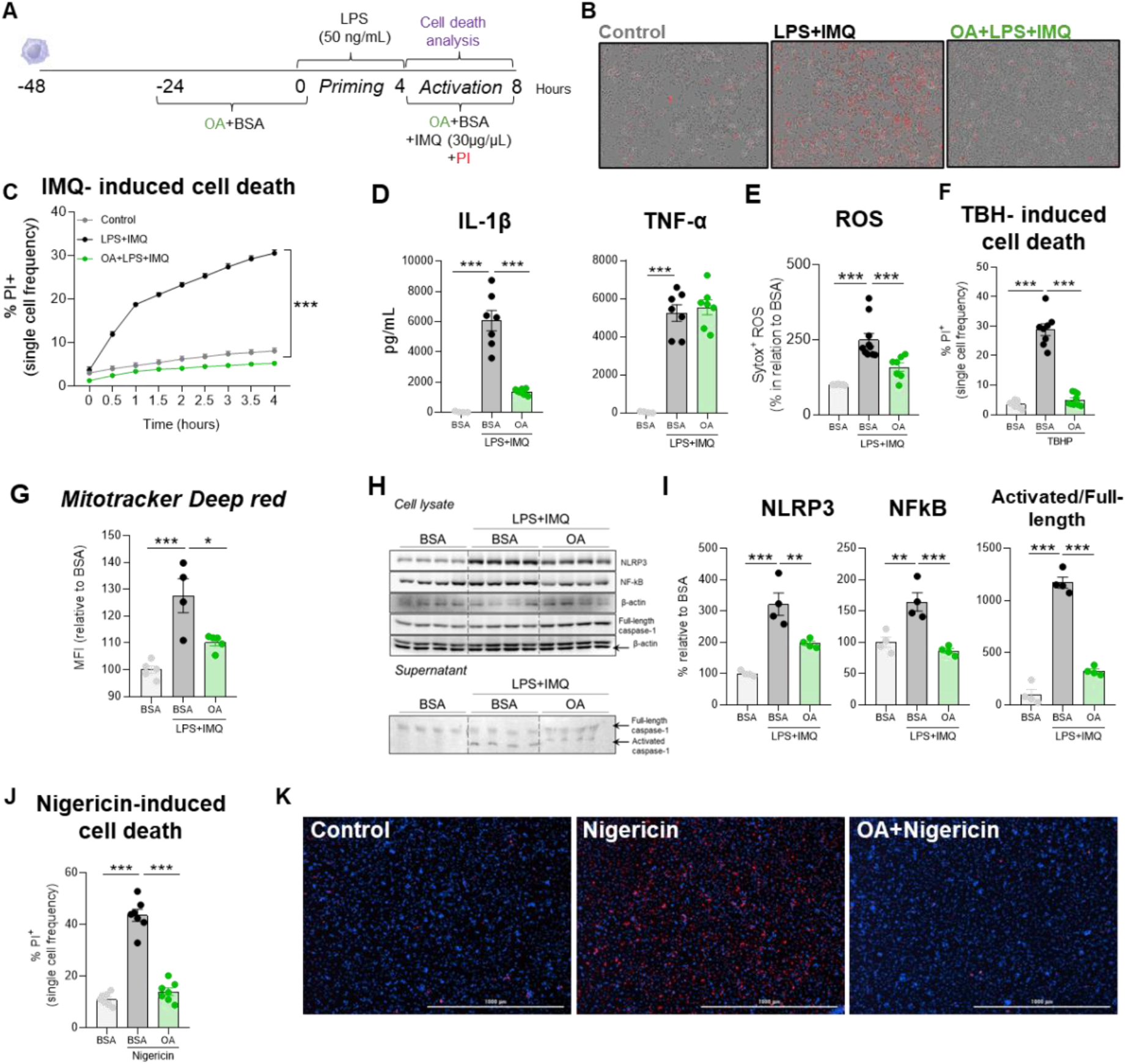
Oleic acid protects macrophages from imiquimod-, TBHP-, and nigericin-induced cell death in vitro. **(A)** Experimental design: dermal macrophages were cultured, primed with LPS for 4 h, and subsequently stimulated with imiquimod. Cell death, cytokine secretion and ROS production were assessed after 4 h of activation. **(B)** Representative images of cell cultures after 4 hours, showing PI staining (red). **(C)** PI incorporation over 4 hours. **(D)** IL-1β and TNF-α concentrations in cell culture supernatants. **(E)** Quantification of ROS production. **(F)** Cell death following 8 hours of treatment with the ROS inducer TBHP. **(G)** MFI of MitoTracker Deep Red, representing mitochondrial membrane potential. **(H)** Protein expression of NLRP3, NF-κB, and cleaved caspase-1 in dermal macrophages lineage, assessed by Western blotting. **(I)** Densitometric quantification of protein expression. **(J)** PI incorporation in BMDMs after 4 hours of stimulation with the specific NLRP3 activator nigericin (5 µM). **(K)** Representative images of cell cultures after 4 hours, showing PI staining (red). Values are expressed as mean ± SEM. (*) p≤0.05; (**) p≤0.01; (***) p≤0.001 by Two-Way ANOVA (C) or one-way ANOVA followed by Dunnett’s post hoc test (D, F, G, I and J).

Oleic acid also limited imiquimod-induced oxidative stress by preventing the rise of reactive oxygen species (ROS) in DMLs **(Figure 5E)** and protected cells from death induced by the direct oxidative stressor TBHP **(Figure 5F)**, indicating a broader protective effect against oxidative damage. Consistent with this, oleic acid increased mitochondrial content and reversed imiquimod-induced mitochondrial hyperpolarization, restoring mitochondrial membrane potential and redox homeostasis **(Figure 5G)**.

At the molecular level, oleic acid reduced NLRP3 and NF-κB expression and attenuated caspase-1 cleavage compared with imiquimod alone **(Figure 5H-I)**, indicating suppression of NLRP3 inflammasome activation. To test whether this reflected a direct effect on the inflammasome itself, we activated NLRP3 directly with nigericin: oleic acid also protected macrophages from nigericin-induced death **(Figure 5J-K)**, confirming that it acts at the level of the inflammasome rather than through an upstream signal.

Finally, we asked whether this protective mechanism depended on TLR7, the receptor through which imiquimod is classically reported to be sensed (37). TLR7 activation alone, using the agonist R848, was not sufficient to induce macrophage death, indicating that TLR7 activation alone is insufficient to account for the cytotoxic response induced by imiquimod **(Supplementary Figure 5A).** Oleic acid also inhibited the increase of IL-1β and TNF-α downstream of TLR7 activation **(Supplementary Figure 5B).** Moreover, blocking TLR7 with the inhibitor M5049 neither prevented imiquimod-induced death nor removed the protective effect of oleic acid on cell death and IL-1β production **(Supplementary Figure 5C-D).** Together, these results indicate that oleic acid’s protective effect is independent of TLR7.

Our findings demonstrate that oleic acid protects macrophages from imiquimod-induced death by limiting oxidative stress and suppressing NLRP3 inflammasome activation and caspase-1 cleavage. This indicates that inflammasome inhibition, rather than modulation of upstream TLR signaling, as the mechanism underlying oleic acid’s protective effect on dermal macrophage survival.

Overall, oral oleic acid supplementation limits this inflammatory pathway, disrupting the self-amplifying cycle linking macrophage death to psoriatic inflammation, thereby preserving the resident macrophage compartment and attenuating disease severity **(Figure 6).**

**Figure 6.**
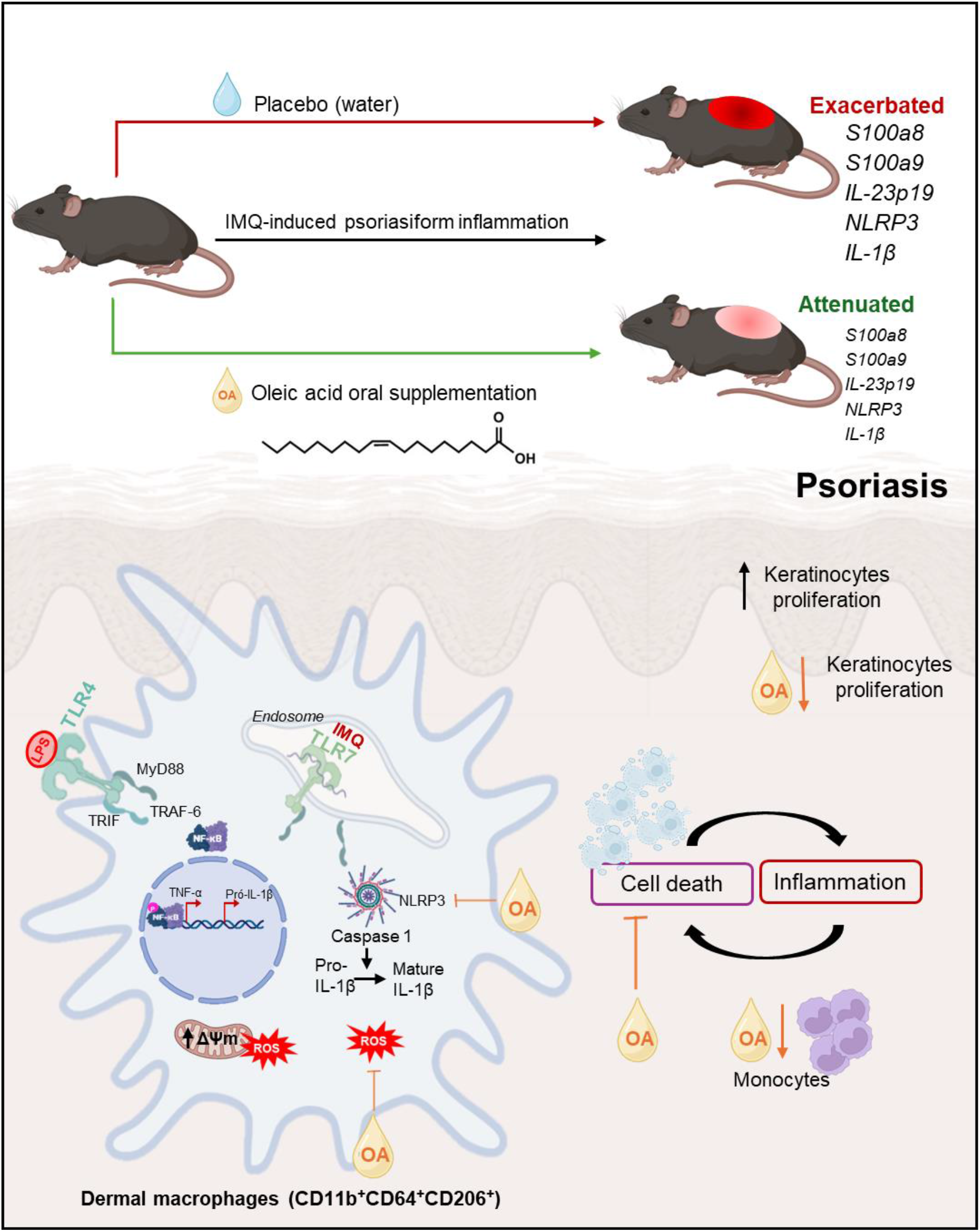
Oleic acid attenuates imiquimod-induced psoriasiform inflammation by protecting dermal macrophages. Schematic summary of the proposed mechanism. Imiquimod promotes inflammatory signaling, ROS production, NLRP3 inflammasome activation, caspase-1 cleavage, and macrophage death, contributing to sustained skin inflammation, keratinocyte proliferation, and recruitment of monocyte-derived cells. Oral oleic acid supplementation attenuates these responses, reduces inflammatory mediators and macrophage death, and preserves the resident dermal macrophage compartment, thereby mitigating psoriasiform inflammation.

## DISCUSSION

Here, we show that the survival of resident dermal macrophages is a key determinant of disease severity in psoriasis, and that dietary oleic acid limits psoriasiform inflammation by directly protecting these cells from inflammasome-driven death. Mechanistically, oleic acid suppressed NF-κB/NLRP3/caspase-1/IL-1β signaling, preserved mitochondrial homeostasis, and limited oxidative stress in macrophages, independently of TLR4- or TLR7-dependent priming. Together, these findings position resident dermal macrophage survival as a therapeutic node in psoriasis and identify dietary oleic acid as a mechanism-based strategy to complement current anti-inflammatory therapies.

Resident dermal macrophages support tissue homeostasis through debris clearance, inflammation resolution, and repair (46,47). Imiquimod markedly reduced this population, with the remaining cells progressively replenished by monocyte-derived inflammatory macrophages. This loss reflected NLRP3/caspase-1-dependent inflammatory cell death, consistent with the established role of inflammasome activation, IL-1β maturation, and the IL-23/IL-17 axis in psoriasis pathogenesis (10–12,48,49), and with reports of caspase-1 activation in patient lesions (49). Pharmacological inhibition of caspase-1 and adoptive transfer of dermal macrophages each attenuated disease in our model, thereby directly linking macrophage death to psoriasiform inflammation.

This cell death coincided with suppressed expression of stearoyl-CoA desaturase (SCD), the rate-limiting enzyme for endogenous oleic acid synthesis (50), in psoriatic skin from both patients and mice, alongside reduced serum oleic acid. This observation suggests a functional link between disrupted lipid metabolism and cutaneous inflammation, distinct from the elevated oleic acid reported in resolved psoriatic lesions (51), and consistent with lipid availability tracking disease activity rather than resolution alone. Normalizing circulating oleic acid levels through oral supplementation inhibited resident dermal macrophage death, potentially by limiting pyroptosis, thereby suppressing inflammatory activation and promoting macrophage survival, maintaining a transcriptional profile similar to the one encountered in healthy skin.

This macrophage-protective effect translated into a clear reduction in disease severity: oral oleic acid attenuated skin thickening and epidermal hyperplasia, blocked upregulation of *S100a8*, *S100a9*, and *Il23p19*, key amplifiers of the IL-23/IL-17 axis (35,36,52), and preserved resident dermal macrophages independently of changes in body weight. These effects are consistent with the anti-inflammatory actions of oleic acid reported in other macrophage and allergic inflammation models (53,54), and extend this evidence to a macrophage-intrinsic mechanism in psoriasis.

Consistent with *in vivo* observations, oleic acid protected macrophages from imiquimod-induced death *in vitro* by suppressing NF-κB and NLRP3 expression and caspase-1 cleavage. Oleic acid also limited NLRP3 activation, as shown by its inhibitory effect against nigericin, and this inhibition was accompanied by reduced oxidative stress and normal mitochondrial membrane potential. Notably, inhibition persisted even when oleic acid was supplied only during the activation phase, or independently of TLR7 engagement, and extended to nigericin-induced death, a stimulus that activates NLRP3 independently of upstream pattern-recognition receptor signaling. Together, these findings indicate that oleic acid’s protective effect does not require intact TLR7 signaling, and is instead accompanied by reduced oxidative stress and preserved mitochondrial membrane potential.

The breadth of these effects, spanning oxidative stress, mitochondrial function, and inflammasome signaling, suggests that oleic acid acts through multiple interconnected mechanisms rather than a single pathway. This is consistent with our use of purified oleic acid rather than olive oil, whose complex lipid composition has yielded conflicting effects on psoriasis (55–57). Because psoriasis remains a chronic disease without a definitive cure (58–60), these findings support dietary oleic acid as a complementary therapeutic strategy and provide a rationale for testing lipid-based nutritional interventions in other chronic inflammatory conditions driven by macrophage dysfunction and inflammatory cell death.

Our findings also identify macrophage survival as a druggable node in psoriasiform inflammation, opening avenues for novel therapeutic approaches targeting immunometabolic checkpoints in psoriasis and, potentially, other macrophage-dependent inflammatory skin diseases.

## METHODS

### Datasets

To gain insight into the molecular mechanisms underlying psoriasis and to identify non-canonical pathways associated with disease pathogenesis, we analyzed the transcriptomic profile of lesional skin from patients with psoriasis and compared it with healthy skin using the publicly available dataset GSE54456 from the National Center for Biotechnology Information (NCBI) Gene Expression Omnibus (GEO).

Differentially expressed genes (DEGs) were identified using GEO2R, an online interactive tool that compares experimental groups based on the original GEO submission data using the GEOquery and limma packages from the Bioconductor project. Volcano plots were generated using GraphPad Prism version 8.0.1.

Functional enrichment analysis of DEGs was performed using the Metascape online platform (61). Genes with a log₂ fold change (log₂FC) ≥ 1.3 were considered upregulated, whereas genes with a log₂FC ≤ −1.3 were considered downregulated. Only genes with an adjusted *P* value < 0.05 were included in downstream analyses. For visualization purposes, enrichment significance was expressed as −log₁₀(*P*), and pathways with −log₁₀(*P*) > 1.3 were considered significantly enriched.

Heatmaps were generated using Morpheus (Broad Institute; https://software.broadinstitute.org/morpheus), and Venn diagrams were constructed using InteractiVenn (62).

### Mice

All animal procedures were approved by the Animal Use Ethics Committee (CEUA) of the University of Campinas (UNICAMP) (protocol no. 5562-1/2020) and by the Institutional Animal Care and Use Committee (IACUC) of MGH, an Association for Assessment and Accreditation of Laboratory Animal Care (AAALAC) International-accredited facility, under Animal Study Protocol 2021N000252. Male and female C57BL/6 mice aged 8–12 weeks were obtained from the Multidisciplinary Center for Biological Investigation (CEMIB, UNICAMP) and from The Jackson Laboratory (USA).

Animals were randomly assigned to experimental groups and housed in ventilated cages (21 × 32 × 20 cm; H × L × W), maintained on a 12-h light/dark cycle at 23 ± 2°C, and provided with standard chow and water *ad libitum*. All experiments were conducted in accordance with the ethical guidelines for animal research established by the Brazilian National Council for the Control of Animal Experimentation (CONCEA).

Details of the experimental design, including sample sizes and treatment regimens, are provided in the corresponding figure legends.

### Imiquimod-Induced Skin Inflammation Model

To induce psoriasis-like skin inflammation, the dorsal skin of mice was shaved using disposable razors (Wilkinson Sword) one day before the initiation of the protocol, as previously described (63). Mice were then anesthetized with isoflurane delivered through a low-flow inhalation anaesthesia system (Bonther^®^, infusion rate of 1.5% and airflow of 0.3 L/min, according to the manufacturer’s instructions). Subsequently, 60 mg of commercially available cream containing 5% imiquimod (Ixium^®^) was topically applied daily to a 4 × 2 cm area of the dorsal skin for 2 or 4 consecutive days. For characterization of the psoriasis-like inflammation model, mice were euthanized on days 1, 3, and 5 after the initiation of treatment.

Body weight was recorded daily throughout the experiments. Dorsal skin thickness was measured at five distinct sites using a digital caliper (Digimess^®^). In addition, the dorsal skin was photographed daily using a digital camera by the same investigator to monitor disease progression.

In experiment with caspase-1 inhibition, 8 mg/kg of VX765 (S2228, Selleck Chemicals). was applied on the dorsal skin.

### Serum Lipid Composition Analysis by Gas Chromatography–Mass Spectrometry (GC–MS)

To determine the serum lipid composition, blood samples were collected by cardiac puncture and allowed to clot at room temperature for 1 h. Samples were then centrifuged at 3,000 rpm for 15 min at 25°C. Serum was collected and immediately stored at −80°C until analysis.

The GC–MS methodology was adapted from Shirai et al. (2005). Briefly, 1 mL of methanolic sodium hydroxide (NaOH), 150 μL of serum, and 25 μL of pentadecanoic acid internal standard solution (1 mg/mL in isooctane) were mixed and vortexed. Samples were heated for 10 min and subsequently cooled in cold water. Next, 2 mL of boron trifluoride (BF₃) esterification reagent was added, and samples were vortexed, heated for 5 min, and cooled again. Subsequently, 2 mL of isooctane was added and mixed by vortexing, followed by the addition of 5 mL of saturated sodium chloride (NaCl) solution. After phase separation, the organic phase was collected using a glass Pasteur pipette.

Samples were diluted 1:2 in isooctane and analyzed using a GCMS-QP2010 Ultra gas chromatograph coupled to a mass spectrometer (Shimadzu®, Kyoto, Japan) operating with a split ratio of 1:10.

Fatty acids were identified by comparing their retention times with those of a commercial fatty acid methyl ester standard mixture (FAME Mix, 37 Components; Supelco–Sigma®, #CRM47885). Fatty acids not represented in the FAME 37 standard were identified based on spectral matching with the GCMS Real Time Analysis® mass spectral library, using a similarity threshold of ≥80%. Quantification was performed by comparing the integrated peak area of each chromatographic peak with that of the internal standard.

### Oleic Acid Administration

Oleic acid (O1008, Sigma-Aldrich®) was administered orally by gavage at doses of 1.76 g/kg body weight (corresponding to 50 μL), concomitantly with imiquimod (imiquimod)-induced psoriasis-like skin inflammation for 2 or 4 consecutive days. Control mice received water as a vehicle control, as previously described (26).

### Histological Analysis of Dorsal Skin

Skin samples from the dorsal region were collected and fixed in 4% formaldehyde prepared in 0.1 M phosphate buffer (pH 7.4) for 24 h at 4°C. Samples were subsequently dehydrated through a graded ethanol series (50–100%), cleared in xylene, and embedded in paraffin (Paraplast®, P3558, Sigma-Aldrich®). Tissue orientation was standardized to obtain transverse sections perpendicular to the skin surface, encompassing the epidermis, dermis, panniculus carnosus, and dermal white adipose tissue.

Paraffin sections (5 μm) were prepared, rehydrated through a descending ethanol series (100–70%), and stained with hematoxylin and eosin (H&E) for evaluation of psoriasis-associated histopathological changes. Images were acquired using an Eclipse 800 photomicroscope (Nikon, Japan) equipped with a CoolSNAP digital camera (Media Cybernetics, USA).

Quantitative analysis of epidermal area was performed using QuPath software version 0.5.1. Three representative images per histological section were acquired using a 20× objective lens (200× total magnification). The epidermal area was manually delineated and quantified, and results were expressed as relative units normalized to the control group.

### Gene expression analysis

Mice were euthanized, and skin punch biopsies (30 mg) were collected and immediately submerged in RNAlater^™^ (76106, Qiagen^®^) according to the manufacturer’s instructions. Biopsies were stored at 4°C until RNA could be extracted. Biopsies were later homogenized in 1mL ice cold TRIzol^TM^ (15596018, Thermo Fisher Scientific^®^) using a Kinematica Polytron^®^ PT2100 tissue homogenizer, and whole RNA was extracted using the Kit RNAeasy mini (74104, Qiagen^®^) according to the manufacturer’s instructions. RNA quality was assessed by the A260/A280 absorbance ratio prior to quantitative real-time PCR (qRT-PCR) analysis. Only samples with A260/A280 values between 1.8 and 2.0 were included, whereas samples outside this range were excluded. cDNA was synthesized from 2 μg of total RNA using the High-Capacity cDNA Reverse Transcription Kit (4368814, Applied Biosystems^®^) in a final reaction volume of 20 μL, according to the manufacturer’s instructions. The reactions were performed using iTaq™ Universal SYBR® Green Supermix (1725121, Bio-Rad^®^) following the manufacturer’s recommended protocol. The primer sequences used in this study are listed in Table 1.

**Table 1.**
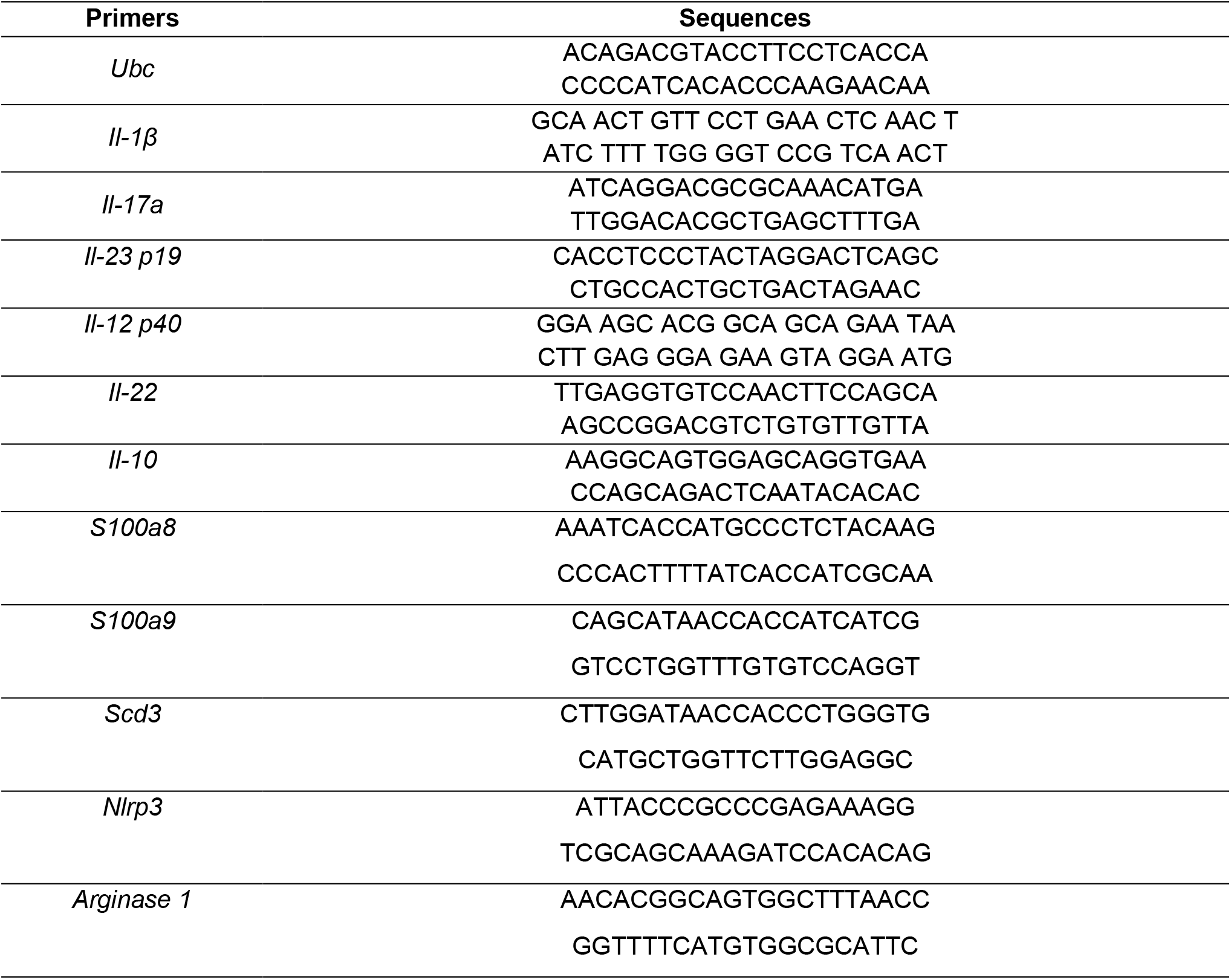
Primers used for quantitative PCR. Sense and antisense primer sequences.

### Skin RNAsequencing

Total RNA was extracted from skin samples following the standard RNA isolation protocol described above and RNA integrity was assessed using the Bioanalyzer 2100 system (Agilent Technologies), and only samples with an RNA Integrity Number (RIN) ≥ 8.6 were considered suitable for downstream analysis. Qualified samples were submitted to Novogene Corporation Inc. (Sacramento, CA, USA) for RNA sequencing.

Sequencing libraries were prepared from poly(A)-enriched mRNA using the NEBNext® Ultra™ RNA Library Prep Kit (New England Biolabs) and sequenced on the Illumina NovaSeq 6000 platform with 150-bp paired-end reads, generating an average of 20 million read pairs per sample.

Raw sequencing data (FASTQ files) underwent quality control, including adapter trimming, removal of low-quality reads, and contaminant filtering using FastQC and Trimmomatic. Filtered reads were aligned to the mouse reference genome (Mus musculus, GRCm39) using HISAT2, and gene expression quantification was performed with featureCounts.

Differential gene expression analysis was conducted using the DESeq2 package. Genes with an absolute log2 fold change (|log2FC|) ≥ 1.3 and p < 0.05 were considered differentially expressed. Functional enrichment analysis of differentially expressed genes was performed using the online platform Metascape (61). For visualization purposes, p-values were transformed to −log10(p), and pathways or biological processes with −log10(p) > 1.3 were considered significantly enriched.

Heatmaps were generated using the Morpheus software, while Venn diagrams were created using the InteractiVenn tool (62). mRNA sequencing will be deposited at NCBI.

### Skin Immunofluorescence

Skin samples were collected and fixed in 4% paraformaldehyde (PFA) for 24h at 4°C. Tissues were then cryoprotected by sequential incubation in 15% and 30% sucrose solutions, each for 24h at 4°C. Subsequently, samples were oriented transversely, embedded in Tissue-Tek® O.C.T. Compound, and rapidly frozen in isopentane pre-cooled in liquid nitrogen. Samples were stored at −80°C until sectioning. Cryosections (10 μm thickness) were obtained using a cryostat at −20°C.

For analysis of keratinocyte proliferation, immunofluorescence staining for Ki67 and keratin 14 (K14) was performed. Cryosections were fixed in acetone for 10 min at −20°C, followed by permeabilization with 0.5% Triton X-100 and SDS solution for 30 min at 4°C. Sections were then blocked with 5% bovine serum albumin (BSA) for 1h at 4°C to reduce non-specific binding.

Samples were incubated overnight at 4°C with a conjugated anti-Ki67 monoclonal antibody (clone SolA15, eFluor 570, eBioscience™, Cat. No. 41-5698-82, 1:100 dilution) and an anti-K14 monoclonal antibody (NBP2-67585, rabbit, 1:200 dilution). The following day, slides were washed with PBS and incubated for 90 min at 4°C with a secondary antibody, Alexa Fluor® 647 donkey anti-rabbit IgG (clone Poly4064, BioLegend, Cat. No. 406414, 1:500 dilution). Nuclei were counterstained using Vectashield H-1000 mounting medium containing DAPI (Vector Laboratories, Cat. No. H-1200-10).

For specificity controls, selected sections were processed without primary antibodies. Images were acquired using a Leica DMi8 inverted microscope equipped with a 40× water-immersion objective (400× total magnification, considering a 10× eyepiece). Ki67⁺ keratinocytes were quantified in three fields per sample using ImageJ software, covering a total analysed area of 70 mm² per sample.

### Flow Cytometry of Skin Samples

The skin flow cytometry protocol was adapted from previously described methods in the literature (Lou; Sun; Wang, 2020; Sakamoto et al., 2022; Willenborg et al., 2022). Skin samples (approximately 300 mg) were collected on day 5 after the initiation of the experimental protocol. Epidermis and dermis were separated by incubating the tissue in Dispase I (LS02100, Worthington) for 15 min at 37°C under constant agitation (200 rpm), followed by mechanical dissociation of the layers using curved forceps.

After separation, dermal tissue was minced with scissors and digested with DNase I from bovine pancreas (200 μg/mL; DN25, Sigma-Aldrich®) and Liberase (30 μg/mL; 5401119001, Sigma-Aldrich®) for 60 min at 37°C under agitation (200 rpm). At the end of digestion, dermal suspensions were washed and filtered through a 100 μm cell strainer to obtain single-cell suspensions.

Cells were incubated with anti-CD16/CD32 blocking antibody (InVivoMAb anti-mouse CD16/CD32, clone 2.4G2, Bio X Cell; 2 μg/mL) for 15 min to reduce non-specific binding. Subsequently, cells were stained for 1h at 4°C in the dark with fluorochrome-conjugated antibodies (Table 2). Samples were then washed and resuspended in DAPI solution (250 ng/mL; D9542, Sigma-Aldrich®) for viability assessment.

**Table 2.**
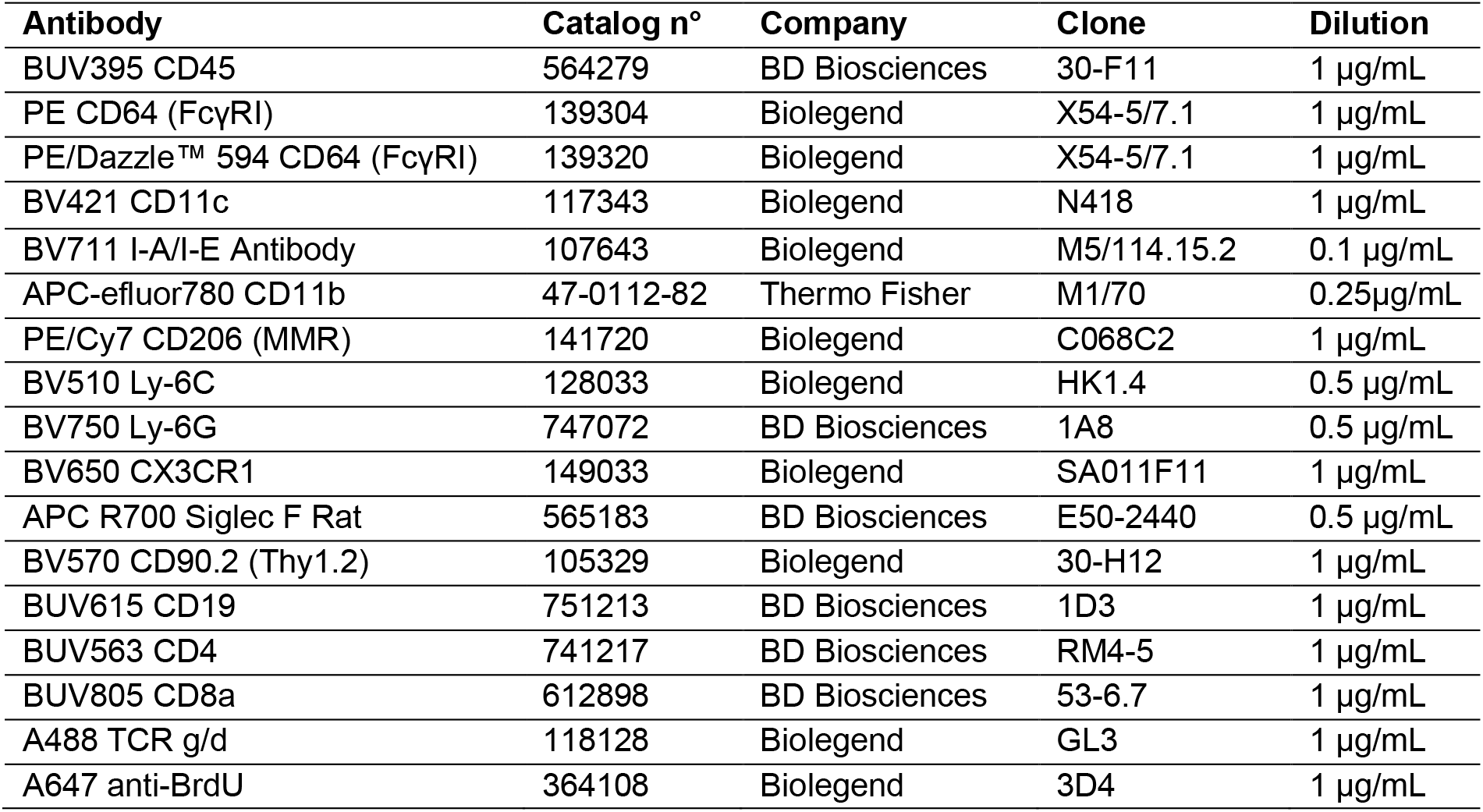
Fluorochrome-conjugated antibodies were used for flow cytometry.

| **Antibody** | **Catalog n°** | **Company** | **Clone** | **Dilution** |
|---|---|---|---|---|
| BUV395 CD45 | 564279 | BD Biosciences | 30-F11 | 1 µg/mL |
| PE CD64 (FcγRI) | 139304 | Biolegend | X54-5/7.1 | 1 µg/mL |
| PE/Dazzle™ 594 CD64 (FcγRI) | 139320 | Biolegend | X54-5/7.1 | 1 µg/mL |
| BV421 CD11c | 117343 | Biolegend | N418 | 1 µg/mL |
| BV711 I-A/I-E Antibody | 107643 | Biolegend | M5/114.15.2 | 0.1 µg/mL |
| APC-efluor780 CD11b | 47-0112-82 | Thermo Fisher | M1/70 | 0.25µg/mL |
| PE/Cy7 CD206 (MMR) | 141720 | Biolegend | C068C2 | 1 µg/mL |
| BV510 Ly-6C | 128033 | Biolegend | HK1.4 | 0.5 µg/mL |
| BV750 Ly-6G | 747072 | BD Biosciences | 1A8 | 0.5 µg/mL |
| BV650 CX3CR1 | 149033 | Biolegend | SA011F11 | 1 µg/mL |
| APC R700 Siglec F Rat | 565183 | BD Biosciences | E50-2440 | 0.5 µg/mL |
| BV570 CD90.2 (Thy1.2) | 105329 | Biolegend | 30-H12 | 1 µg/mL |
| BUV615 CD19 | 751213 | BD Biosciences | 1D3 | 1 µg/mL |
| BUV563 CD4 | 741217 | BD Biosciences | RM4-5 | 1 µg/mL |
| BUV805 CD8a | 612898 | BD Biosciences | 53-6.7 | 1 µg/mL |
| A488 TCR g/d | 118128 | Biolegend | GL3 | 1 µg/mL |
| A647 anti-BrdU | 364108 | Biolegend | 3D4 | 1 µg/mL |

Data acquisition was performed using a BD FACSymphony™ A5 flow cytometer (BD Biosciences, Maryland, USA), and data analysis was carried out using FlowJo™ software v10 (BD Biosciences, Maryland, USA).

The gating strategy used for analysis is shown in **Supplementary figure 6**.

### In vivo proliferation analysis of skin myeloid cells

For in vivo proliferation assessment, bromodeoxyuridine (BrdU; 40 mg/kg; 100 μL; B5002-1G, Sigma-Aldrich®) was administered intraperitoneally every 12 h on days 1–2 or 3–4 of the experimental protocol. Dorsal skin was collected on day 3 or day 5 for preparation of single-cell suspensions, as described before.

Cells were first stained with Live/Dead viability dye (L34962, Thermo Fisher Scientific®) for 30 min at 4°C. Suspensions were then incubated with anti-CD16/CD32 blocking antibody for 15 min, followed by staining with extracellular antibodies (Table 2) for 1h at 4°C in the dark. Samples were subsequently fixed and permeabilized using the BD Cytofix/Cytoperm™ Fixation/Permeabilization Kit (BDB554714, Fisher Scientific) together with BD Permeabilization Buffer Plus (56165, BD Biosciences), according to the manufacturer’s instructions.

To expose incorporated BrdU, cells were treated with DNase I (800 μg/mL) and incubated for 1 h at 37°C. Intracellular staining was then performed using an anti-BrdU Alexa Fluor® 647 antibody (Table 2), with overnight incubation at 4°C.

Data acquisition was performed using a BD FACSymphony™ A5 flow cytometer (BD Biosciences, Maryland, USA), and data analysis was carried out using FlowJo™ v10 software (BD Biosciences, Maryland, USA). The percentage of BrdU-positive myeloid cells was quantified.

### Dermal macrophage restoration

To restore the dermal macrophage population, mice received subcutaneous injections of 2.5 × 10⁵ dermal macrophages per mouse (64), suspended in PBS, concomitantly with topical imiquimod treatment. Control mice received an equivalent volume of PBS under the same experimental conditions.

### Western blotting of skin

Skin samples (100 mg) were homogenized in RIPA lysis buffer supplemented with protease and phosphatase inhibitors (Santa Cruz Biotechnology, sc-24948), and centrifuged at 12,000 rpm for 10 min at 4°C. Protein concentration was determined using the Pierce™ BCA Protein Assay Kit (23225, Thermo Fisher Scientific®), and samples were adjusted to 1 μg/μL.

Laemmli buffer containing 100 mM DTT was added to the lysates, followed by heating at 95°C for 10 min. Equal amounts of protein (20 μg) were separated by SDS-PAGE and transferred onto nitrocellulose membranes. Membranes were blocked with 5% non-fat milk in TBS-T (0.2% Tween-20) for 60 min at room temperature and incubated overnight at 4°C with primary antibodies.

The following primary antibodies were used: anti-caspase-1 (p20) (1 μg/mL, AG-20B-0042-C100, Adipogen®) and anti-β-actin (rabbit, 0.5 μg/mL, 600-401-886, Rockland). Membranes were washed and incubated with appropriate secondary antibodies for 2 h at room temperature. Protein bands were detected using chemiluminescent substrate and imaged with a G XR5 gel imaging system (Syngene®, Cambridge, UK). Band intensities were quantified using ImageJ software.

### Analysis of the resilience of skin-resident macrophages using an endocytic labelling strategy

Taking advantage of the high endocytic capacity of resident macrophages, dextran–rhodamine labeling was used to track the resilience of these cells throughout the experimental protocol. Dextran–rhodamine 70 kDa (R9379, Sigma-Aldrich®, 5 mg/kg, 50 μL) was administered via the retro-orbital route between days 0 and 1 of the protocol. On day 5 after the beginning of the experiment, skin samples were collected and processed for flow cytometry as described. The percentage of dextran-positive myeloid cells was quantified.

### *In vitro* experiments

#### Dermal Macrophage Cell Culture

Cells were cultured at 37°C/5% CO2 in a humidified incubator unless indicated otherwise. Dermal macrophage cell lines (DMLs) were kindly provided by Prof. Carlos H. Serezani (Vanderbilt University, USA) and generated as previously described (65,66). Immortalized cells were maintained in culture in Dulbecco’s Modified Eagle’s Medium (DMEM) containing 4.5 g/L glucose supplemented with 10% heat-inactivated fetal bovine serum (FBS), 100 units/mL penicillin (15140122), 100 μg/mL amphotericin B (15290026), and 2 mM L-glutamine (all from Gibco Invitrogen, Grand Island, NY, USA). Cultures were passaged every 2 days and used up to passage 10. All cell culture assays were performed in duplicate and independently repeated three times.

#### Generation of Bone Marrow-Derived Macrophages (BMDMs)

Murine bone marrow-derived macrophages (BMDMs) were generated from bone marrow cells isolated from the femurs and tibias of C57BL/6 mice (67). Following red blood cell lysis with ammonium-chloride-potassium (ACK) buffer, cells were cultured at densities appropriate for each experiment in RPMI medium supplemented with 10% FBS and differentiated in the presence of macrophage colony-stimulating factor (M-CSF; 20 ng/mL; PMC2044; Thermo Fisher Scientific) for 5 days.

#### Cell Death Analysis by Propidium Iodide Incorporation

DMLs (1.2 × 10⁴ cells/well) or BMDMs (1.2 × 10⁵ cells/well) were seeded in 96-well plates. After 5 days of differentiation, the cells were primed with 50 ng/mL of LPS from E. coli O111:B4 (IAX100012M001, Adipogen) for 4 h. After priming, the medium was replaced and cells were stimulated with increasing concentrations of imiquimod (tlrl-imqs-1, InvivoGen) (10–45 µg/mL) in the presence of propidium iodide (PI; final dilution 1:300). To evaluate the effects of oleic acid on imiquimod-induced cell death, cells were pretreated with 300 µM oleic acid conjugated to BSA (O3008-5ML, Sigma-Aldrich) for 24 h and subsequently primed with LPS for 4 h. After priming, cells were incubated with OA-BSA, imiquimod (15 or 30 µg/mL; tlrl-imqs-1, InvivoGen), and PI (1:300). Time-lapse imaging was performed using an Incucyte® Live-Cell Analysis System (Sartorius) or a Cytation 5 Cell Imaging Multi-Mode Reader (Agilent/BioTek) maintained at 37°C and 5% CO₂. Brightfield and red fluorescence images were acquired every 30 min or 1 h for 4 h. For Cytation 5 experiments, Hoechst (H3570) was included for nuclear staining. Images were acquired using a 10× Plan Fluor objective, and acquisition parameters were kept constant across experimental groups. Image analysis was performed using Gen5™ software with automated object segmentation. To investigate the mechanisms of imiquimod-induced cell death, cells were treated with the NLRP3 inhibitor MCC950 (1 µM; S7809, Selleck Chemicals), the caspase-1 inhibitor VX765 (25 µM; S2228, Selleck Chemicals). To assess the involvement of TLR7 signaling, cells were stimulated with the TLR7 agonist resiquimod (R848; 30 µg/mL; tlrl-r848-1, InvivoGen) or treated with the selective TLR7/8 inhibitor entospletinib (0.5 µM; M5049, inh-m5049, InvivoGen).

Plates were equilibrated with PI for 30 min prior to image acquisition to optimize exposure, gain, and focus settings.

#### Immunodetection of proteins

For cytokine quantification, cells were seeded in 24-well plates at a density of 2.5 × 10⁵ cells/well. Following differentiation and experimental stimulation, cell-free supernatants were collected, and IL-1β (DY401-05) and TNF-α (DY410-05) concentrations were quantified using DuoSet ELISA kits (R&D Systems, Minneapolis, MN, USA) according to the manufacturer’s instructions.

For immunoblot analysis, cells were seeded in 6-well plates at a density of 1 × 10⁶ cells/well. Following differentiation and experimental stimulation, cell-free supernatants and cell lysates (in RIPA buffer supplemented with protease and phosphatase inhibitors (sc-24948, Santa Cruz Biotechnology) were collected. Lysates were centrifuged (12,000 rpm, 10 min, 4°C) and protein concentrations were determined using the Pierce™ BCA Protein Assay Kit (23225, Thermo Fisher Scientific) and normalized to 1 µg/µL. Samples were mixed with Laemmli buffer containing 100 mM DTT and heated at 95°C for 10 min. Equal amounts of protein (20 µg) were separated by SDS-PAGE and transferred onto nitrocellulose membranes.

Membranes were blocked with 5% non-fat milk in TBS containing 0.2% Tween-20 for 1 h at room temperature and incubated overnight at 4°C with the following primary antibodies: anti-NLRP3/NALP3 (Cryo-2; 1 µg/mL, AG-20B-0006, Adipogen), anti-NF-κB p65 (C-20; 0.1 µg/mL, sc-372, Santa Cruz), and anti-caspase-1 p20 (1 µg/mL, AG-20B-0042-C100, Adipogen).

Membranes were washed and incubated with the appropriate secondary antibodies for 2 h at room temperature. Protein bands were detected using chemiluminescent substrate and imaged using a G XR5 Gel Imaging System (Syngene, Cambridge, UK). Band intensity was quantified using ImageJ software. Lysate proteins were normalized to β-actin (0.2 µg/mL, sc-8432-HRP, Santa Cruz).

#### Intracellular Reactive Oxygen Species (ROS) Measurement by Flow Cytometry

Cells were seeded in 24-well plates at a density of 1 × 10⁵ cells/well. After 24 h, cells were treated with 300 µM OA-BSA for 24 h and then primed with LPS for 4 h. The medium was replaced, and cells were incubated with OA-BSA and 30 µg/mL of imiquimod for 30 min.

Intracellular ROS levels were measured using the CellROX™ Orange Flow Cytometry Assay Kit (C10493, Thermo Fisher Scientific). CellROX Orange was added at a final concentration of 750 nM and incubated for 30 min. Cells were detached using TrypLE Express (12604021, Thermo Fisher Scientific) and resuspended in DAPI-containing solution (150 ng/mL) to exclude dead cells.

Samples were analyzed using a BD FACSymphony™ A5 flow cytometer and FlowJo™ v10 software. The median fluorescence intensity (MFI) of viable cells was used to quantify intracellular ROS production.

### RNA Extraction and Quantitative Real-Time PCR

Cells were seeded in 24-well plates at a density of 2.5 × 10⁵ cells/well. Following 4 h of LPS priming, cells were stimulated with imiquimod for an additional 4 h in the presence or absence of 300 µM OA-BSA. Cells were then lysed in 1 mL of TRIzol reagent, and total RNA was extracted according to the manufacturer’s instructions.

RNA quality was assessed by the A260/A280 absorbance ratio. Complementary DNA (cDNA) was synthesized from 2 µg of RNA using the High-Capacity cDNA Reverse Transcription Kit (4368814, Applied Biosystems) in a final volume of 20 µL. Quantitative real-time PCR was performed using iTaq Universal SYBR Green Supermix (1725121, Bio-Rad) according to the manufacturer’s instructions. Primer sequences are listed in Table 1.

### Statistical Analysis

Results are presented as mean ± standard error of the mean (SEM), and the statistical tests applied are specified in the corresponding figure legends. Statistical analyses were performed using GraphPad Prism 8.0^®^ (GraphPad Software, Inc., San Diego, CA, USA). Differences were considered statistically significant when p < 0.05.

Data normality was assessed using the Shapiro–Wilk test, with p > 0.05 indicating a normal distribution. For normally distributed data, comparisons between two groups were performed using the unpaired Student’s t-test. Comparisons involving more than two groups were analyzed using one-way analysis of variance (ANOVA), whereas experiments involving multiple factors, including time-course analyses, were evaluated using two-way ANOVA. ANOVA analyses were followed by Dunnett’s or Bonferroni’s post hoc tests, as appropriate.

For data that did not meet the assumption of normality, the Mann–Whitney test was used. Correlation analyses were performed using Spearman’s rank correlation coefficient.

## Supporting information

Images of the WB membranes

## Supplementary Figures

**Supplementary Figure 1.**
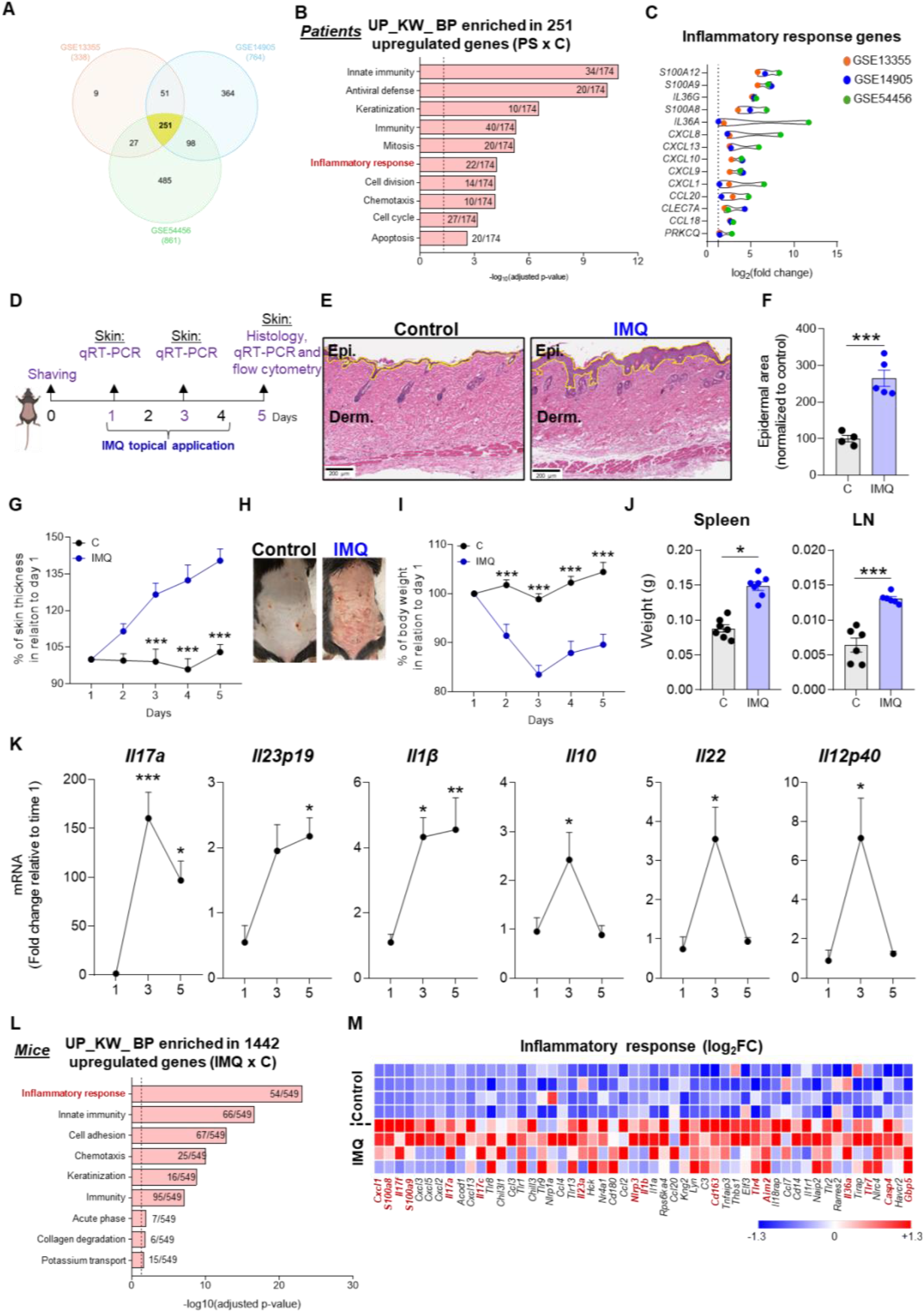
Human and murine psoriasis exhibit conserved inflammatory transcriptional reprogramming. **(A)** Venn diagram showing the overlap of upregulated genes identified across three human psoriasis datasets, considering log_2_(fold change) ≥1.3 or ≤−1.3 and adjusted p-value<0.05. **(B)** Gene ontology (GO) of Biological Processes enriched in 251 commonly upregulated genes in psoriatic versus healthy skin across the three datasets. **(C)** log_2_ fold change of inflammatory response related genes in the three datasets. **(D)** Experimental design: dorsal skin was shaved one day before psoriasis induction, followed by daily topical application of 60 mg of 5% imiquimod cream from day 1 to day 4. Skin thickness and body weight were monitored daily. Skin samples were collected on days 1, 3 and 5 for qRT-PCR analysis and on day 5 for histological analysis. Spleens and draining lymph nodes (LN) were harvested for weight measurement. **(E)** Representative histological images of the skin collected on day 5. Scale bar: 200 µm. **(F)** Quantification of epidermal area relative to the control group. **(G)** Percentage of skin thickness relative to day 1. **(H)** Representative photographs of mouse dorsal skin. **(I)** Percentage change in body weight relative to day 1. **(J)** Spleen and axillary lymph node weights on day 3. **(K)** Skin cytokine gene expression at different time points during imiquimod-induced psoriasis, including *interleukin (Il)-17a*, *Il23p19*, *Il1β*, *Il10*, *Il22*, and *Il12p40*. **(L)** Top upregulated Biological Process identified in the skin following imiquimod treatment. **(M)** Heatmap of genes associated with inflammatory response. In red we highlighted the genes that appeared in the human datasets. Each row represents an individual mouse and each column represents a gene. Values are expressed as mean ± SEM. (*) p≤0.05; (**) p≤0.01; (***) p≤0.001 by unpaired Student’s t-test (F and J) or by two-way ANOVA followed by Bonferroni’s post hoc test (G and I) or by one-way ANOVA followed by Dunnett’s post hoc test (K).

**Supplementary Figure 2.**
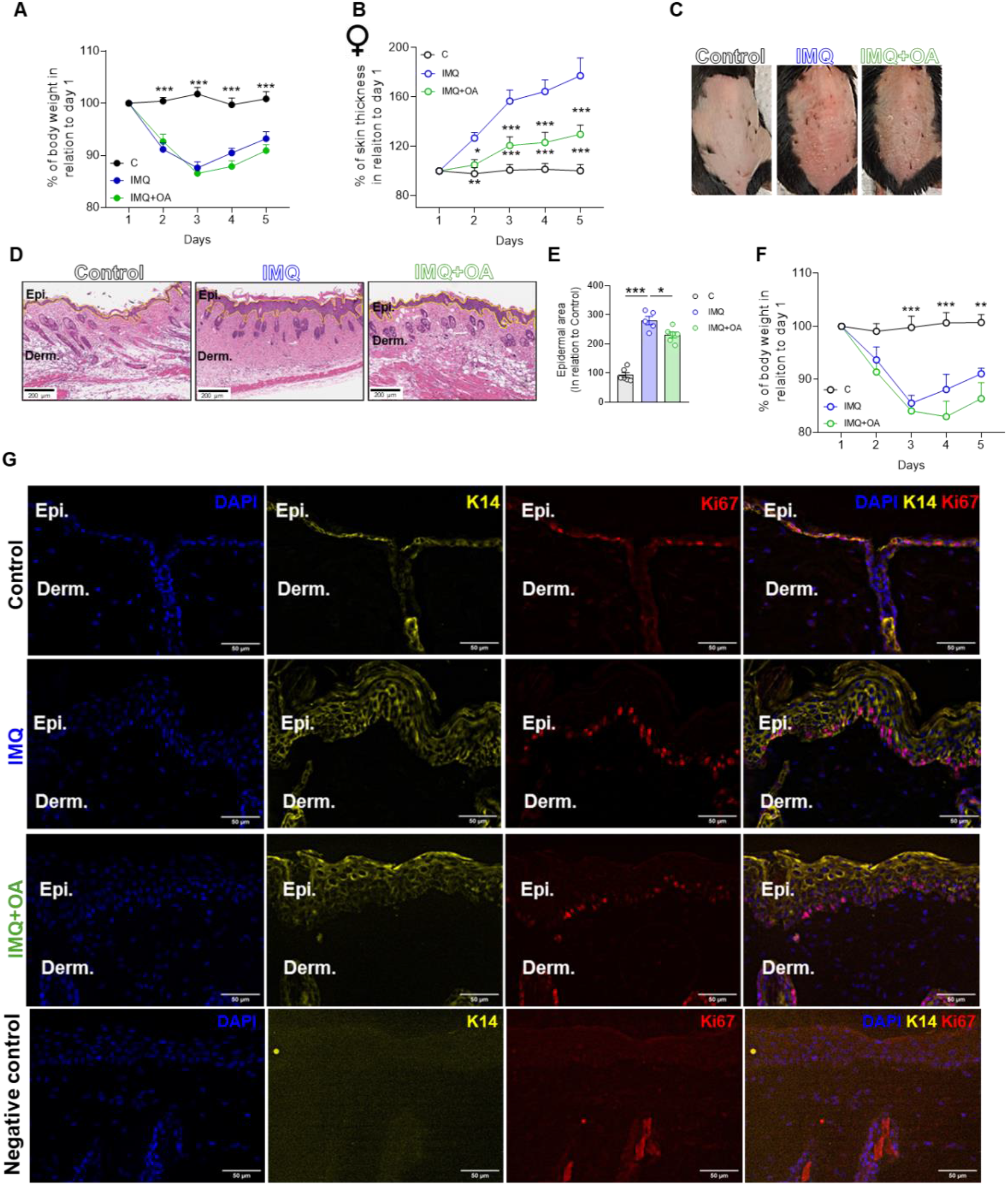
Oral oleic acid supplementation similarly attenuates imiquimod-induced psoriasis in female mice. **(A)** Percentage change of body weight relative to day 1 in male mice. **(B)** Percentage of skin thickness relative to day 1 in female mice. **(C)** Representative photographs of mouse dorsal skin from female mice. **(D)** Representative histological images of skin on day 5 from female mice. Scale bar: 200 µm. **(E)** Quantification of epidermal area relative to the control group. **(F)** Percentage change in body weight relative to day 1 in female mice. **(G)** Representative immunofluorescence images of mouse skin on day 5. Images were acquired using a Leica microscope with a 40× water-immersion objective to detect DAPI (nuclei, blue), K14 (keratinocytes, yellow), and Ki67 (proliferating cells, red). Scale bar: 50 µm. Values are expressed as mean ± SEM. (*) p≤0.05; (**) p≤0.01; (***) p≤0.001 by two-way ANOVA followed by Bonferroni’s post hoc test (A, B and F) or by one-way ANOVA followed by Dunnett’s post hoc test (E).

**Supplementary Figure 3.**
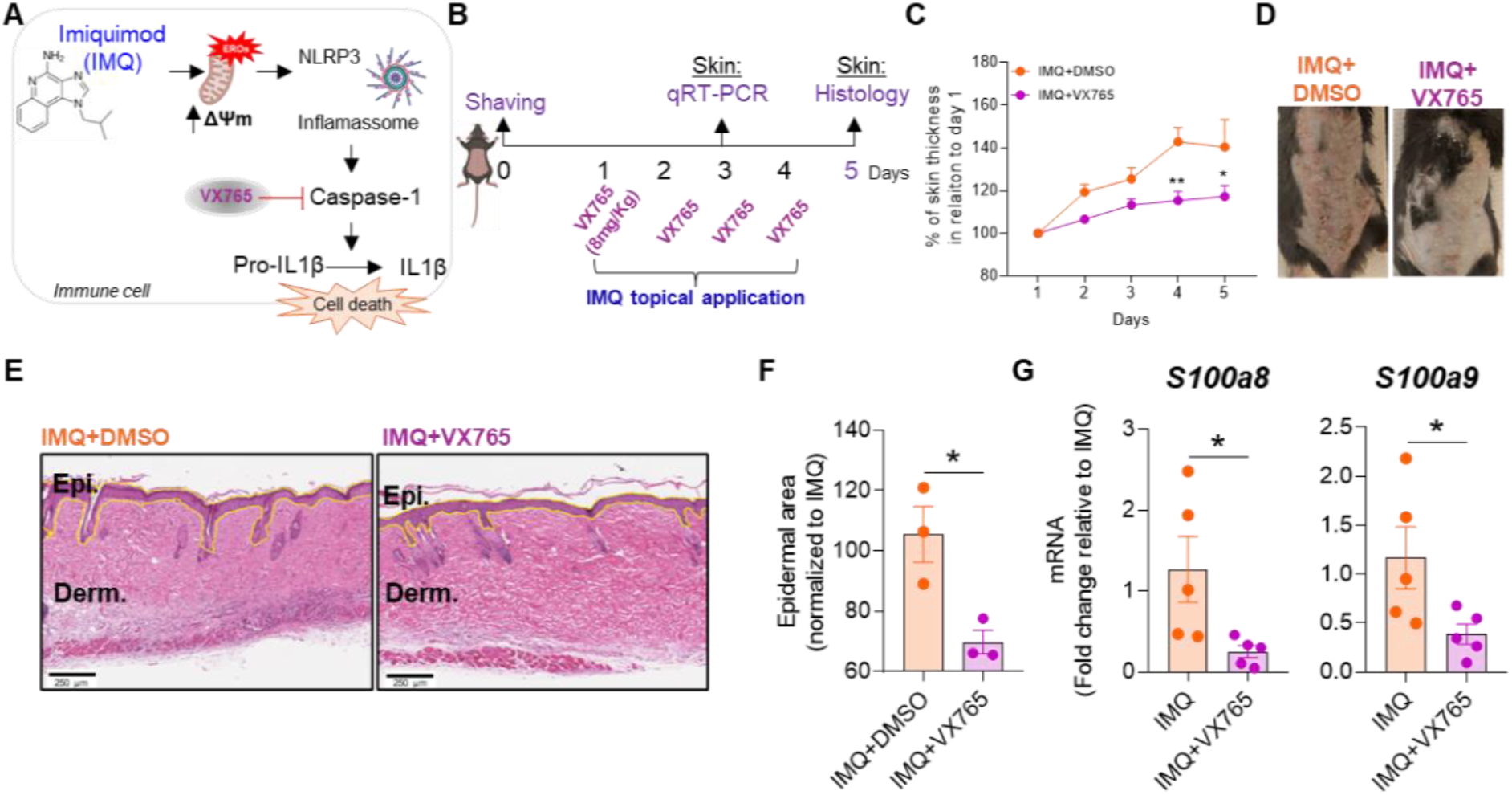
**(A)** Illustration of inflammasome activation and downstream signaling in myeloid cells. **(B)** Experimental design for caspase-1 inhibition during imiquimod-induced psoriasiform inflammation: dorsal skin was shaved one day before disease induction, followed by daily topical application of 60 mg of 5% imiquimod cream from day 1 to day 4. Mice were concom itantly treated topically with VX765 (8 mg/kg). Skin thickness was monitored daily, skin samples were collected on day 3 for qRT-PCR or 5 for histology. **(C)** Percentage of skin thickness relative to day 1. **(D)** Representative photographs of mouse dorsal skin. **(E)** Representative histological images of dorsal skin collected on day 5. Scale bar: 250 µm. **(F)** Quantification of epidermal area relative to the control group. **(G)** Relative gene expression of *S100a8* and *S100a9*, in skin samples collected on day 3. Values are expressed as mean ± SEM. (*) p≤0.05; (**) p≤0.01; (***) p≤0.001 by two-way ANOVA followed by Bonferroni’s post hoc test (C) or by unpaired Student’s t-test (F, G).

**Supplementary Figure 4.**
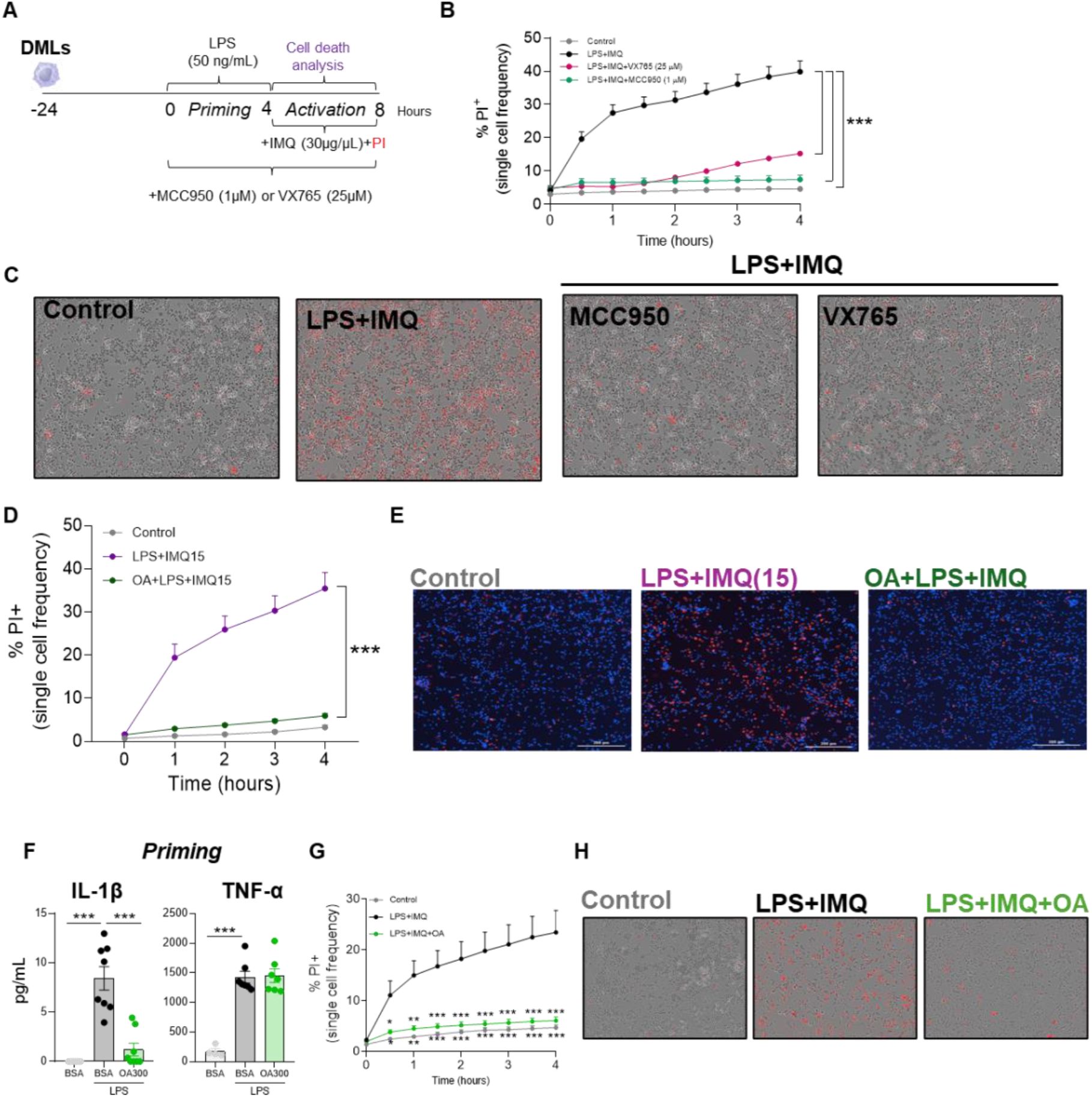
Oleic acid protects macrophages from imiquimod-induced inflammasome-dependent cell death independently of TLR4 priming. **(A)** Experimental design: dermal macrophages were cultured, primed with LPS for 4 h, and subsequently stimulated with imiquimod in the presence or absence of the NLRP3 inhibitor MCC950 or the caspase-1 inhibitor VX765. **(B)** PI incorporation over 4 hours. **(C)** Representative images of PI-positive dermal macrophages after 4 hours with NLRP3- or caspase-1 inhibitor. **(D)** PI incorporation over 4 h in BMDMs pretreated with oleic acid for 24 h and subsequently stimulated with imiquimod (15 µg/mL). **(E)** Representative images of PI-positive cells after 4 h, showing Hoechst (nuclei, blue) and PI staining (red). **(F)** IL-1β and TNF-α concentrations in cell culture supernatant measured by ELISA after 4 hours of LPS priming. **(G)** Macrophages were primed with LPS for 4 hours and subsequently treated with oleic acid (300 µM) and imiquimod. Cell death was evaluated by PI incorporation over 4 hours. **(H)** Representative images of PI-positive cells after 4 hours. Values are expressed as mean ± SEM. (*) p≤0.05; (**) p≤0.01; (***) p≤0.001 by two-way ANOVA followed by Dunnett’s post hoc test (B, D and G) or by one-way ANOVA followed by Dunnett’s post hoc test (F).

**Supplementary Figure 5.**
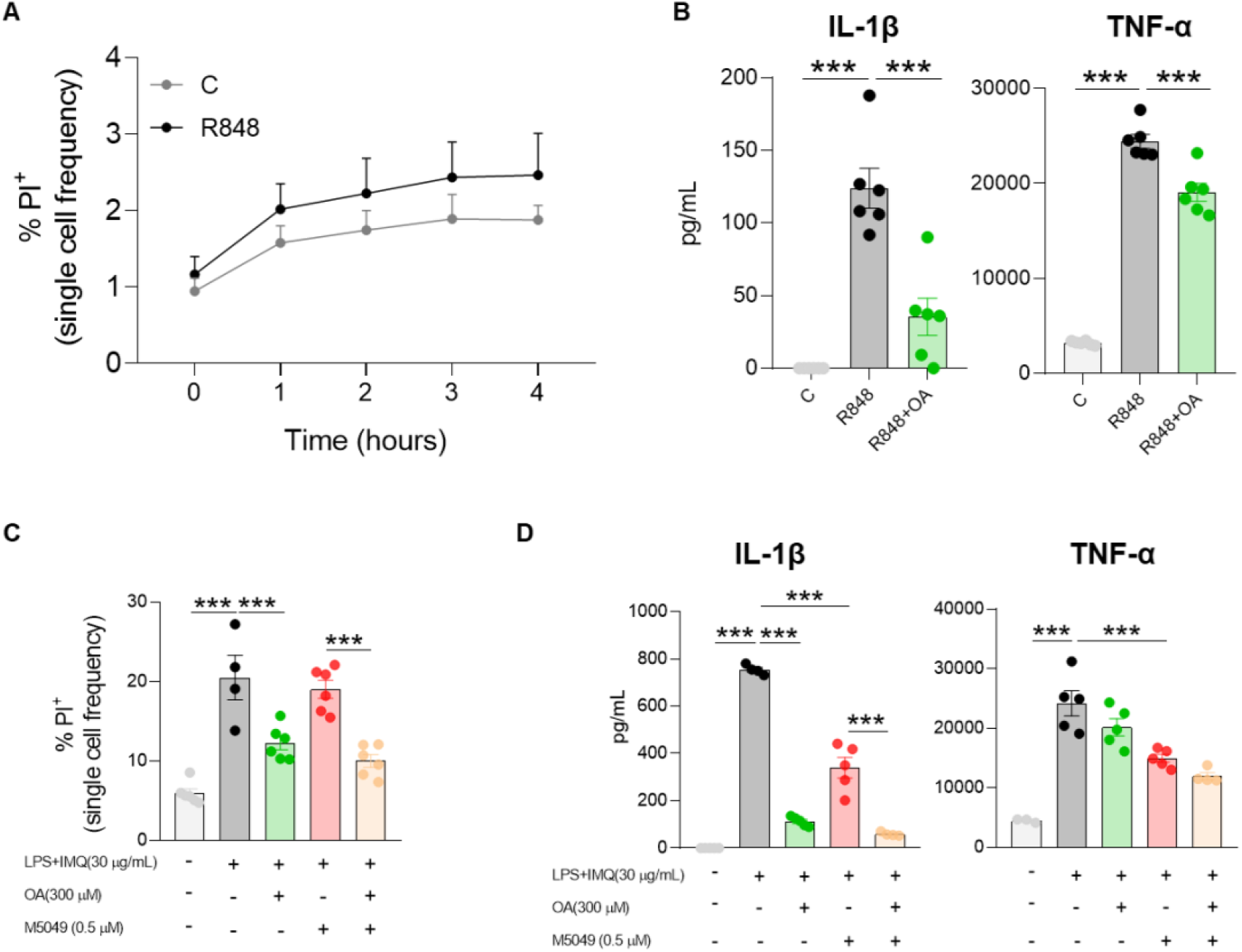
The protective effect of oleic acid (OA) against imiquimod-induced cell death is independent of TLR7. **(A)** Dermal macrophages were stimulated with R848, and cell death was assessed by PI incorporation over 4 hours. **(B)** Dermal macrophages were pretreated for 24 h with 300 µM BSA-conjugated oleic acid and subsequently stimulated with R848 for 8 h. IL-1β and TNF-α concentrations in cell culture supernatants were measured by ELISA after 8 hours of R848 stimulation. **(C)** DLMs were stimulated with LPS and imiquimod in the presence or absence of the TLR7 inhibitor M5049 (0.5 µM), and cell death was assessed by PI incorporation after 8 hours. **(D)** IL-1β and TNF-α concentrations in cell culture supernatants measured by ELISA after 4 hours of imiquimod stimulation. Values are expressed as mean ± SEM. (*) p≤0.05; (**) p≤0.01; (***) p≤0.001 by two-way ANOVA followed by Bonferroni’s post hoc test (A, C and D) or by one-way ANOVA followed by Dunnett’s post hoc test (B).

**Supplementary Figure 6.**
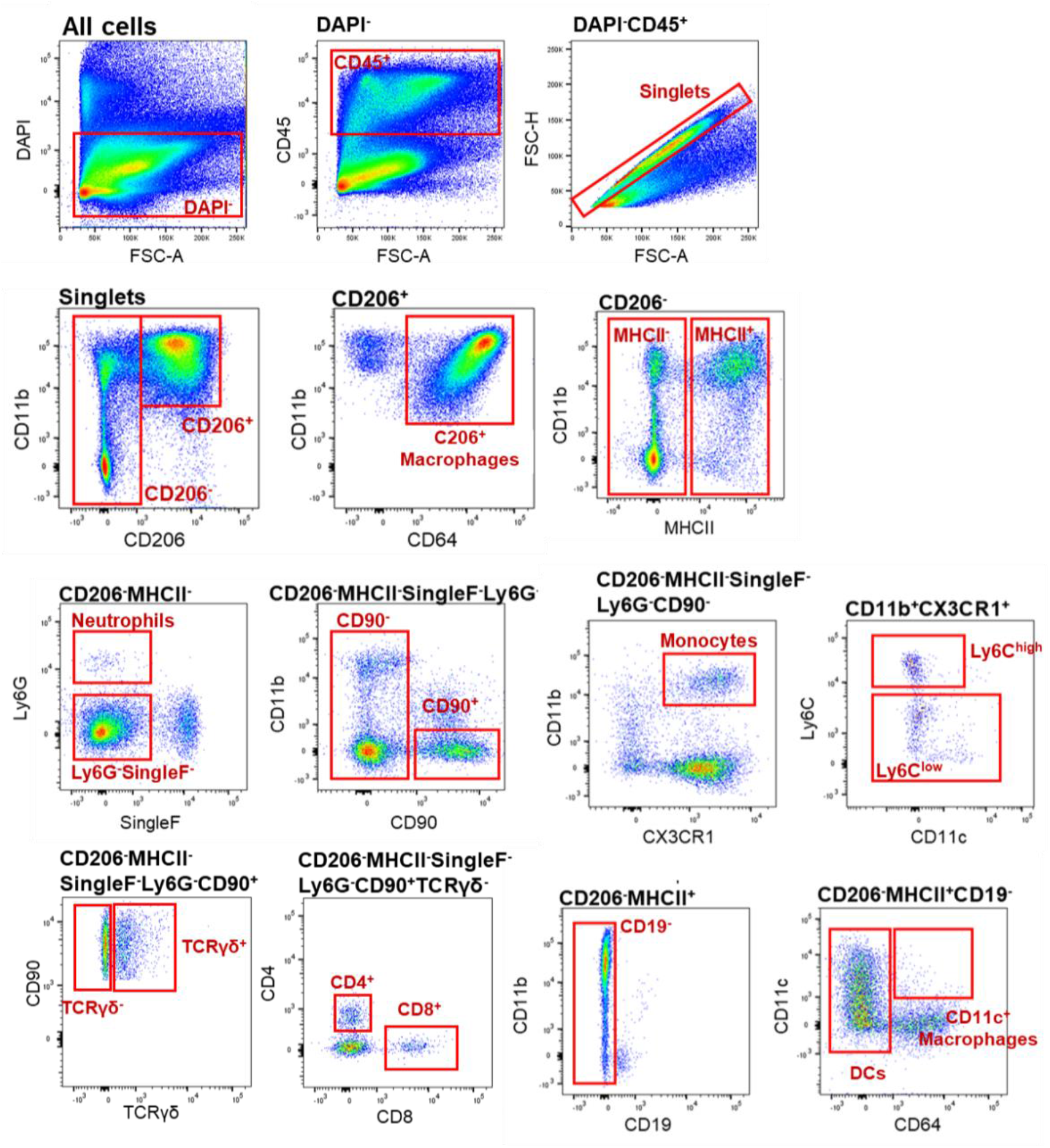
Gating strategy for the characterization of skin immune cells. After exclusion of dead cells (DAPI^+^), we selected the leukocytes (CD45^+^) and excluded the doublets. Myeloid cells were identified based on CD11b and CD206 expression. CD11b^+^CD206^+^ cells were further analysed for CD64 expression, with CD11b^+^CD206^+^CD64^+^ cells defined as resident tissue macrophages. CD206^-^MHCII^-^ cells were analysed for SingleF and Ly6G expression; CD206^-^MHCII^-^Ly6G^+^SingleF^-^ cells were identified as neutrophils. Ly6G^-^SingleF^-^ cells were subdivided according to CD90 expression. CD90^-^CX3CR1^+^ were classified as monocytes and further separated into Ly6C^high^ and Ly6C^low^ subsets. CD206^-^MHCII^+^ cells were analysed for CD19 and CD11c^/^CD64 expression, with CD19⁻CD11c⁺CD64⁺ cells identified as dendritic cells. Lymphoid populations were gated within the CD206^-^ fraction. MHCII^+^CD19^+^ cells were defined as B cells, while CD206^-^MHCII^-^CD90^+^ T cells were subdivided into TCRγδ^+^ and TCRγδ^-^ populations, followed by CD4 and CD8 analysis.

## ACKNOWLEDGEMENTS

We thank the Koch Institute Histology Core at the Massachusetts Institute of Technology (RRID:SCR_019177) for histological sample processing, the members of the Ragon Institute Flow Cytometry Core for expert assistance with flow cytometry, the staff of the Ragon Institute Animal Facility for outstanding animal care and mouse husbandry, the Microscopy Core Facility at the School of Applied Sciences, University of Campinas (FCA–UNICAMP), for technical assistance with microscopy and image acquisition, the VUMC Analytical Services Core for technical support with serum fatty acid analysis, and Prof. Dennys and the members of the Nutritional Genomics Laboratory for providing access to GC-MS instrumentation and technical assistance with fatty acid profiling.

## AUTHOR CONTRIBUTIONS

Conceptualization: B Burger, HM Silva and HG Rodrigues. Experimental design and investigation: B Burger, C Evavold, JCA Filho, P Varga-Weisz, CHC Serezani, MAR Vinolo, HM Silva and HG Rodrigues. Specimen collection: B Burger, RN Sagiorato, IB Baccarin, VR Muñoz, G Cerna, I Andreev, J Zhao, B Sullivan, RAM Bistaffa, VC Gaspar, NB Ramos, LP Pral. Data analysis: B Burger, VR Muñoz, I Andreev, N Araújo, Y Liang, EM Yabuuti, HM Silva and HG Rodrigues. Funding acquisition: B Burger, HM Silva and HG Rodrigues. Manuscript preparation and revision: B Burger, JCA Filho, P Varga-Weisz, CHC Serezani, MAR Vinolo, HM Silva and HG Rodrigues. All authors read and approved the final manuscript.

## ETHICS APPROVAL AND CONSENT TO PARTICIPATE

The mouse maintenance and experiments were performed following the approved protocols by the Animal Use Ethics Committee/UNICAMP (CEUA – 5562-1/2020) and by the Institutional Animal Care and Use Committee (IACUC) of MGH, an Association for Assessment and Accreditation of Laboratory Animal Care (AAALAC) International-accredited facility, under Animal Study Protocol 2021N000252.

The experiments were conducted in accordance with the National Institutes of Health (NIH) Guide for the Care and Use of Laboratory Animals and ARRIVE guideline. The procedures were performed at the Laboratory of Nutrients and Tissue Repair, School of Applied Sciences, UNICAMP, Brazil and at the Animal Facility, Ragon Institute of Mass General Brigham, MIT, and Harvard, Cambridge, MA.

## DATA AVAILABILITY

The data that support the findings of this study are available upon request from the corresponding author. The RNAseq reads will be deposited at NCBI.

