## Supplementary material for "Oleic acid suppresses NLRP3 inflammasome activation to prevent dermal macrophage death and limit imiquimod-induced psoriasis": Images of the WB membranes

### Supplementary file

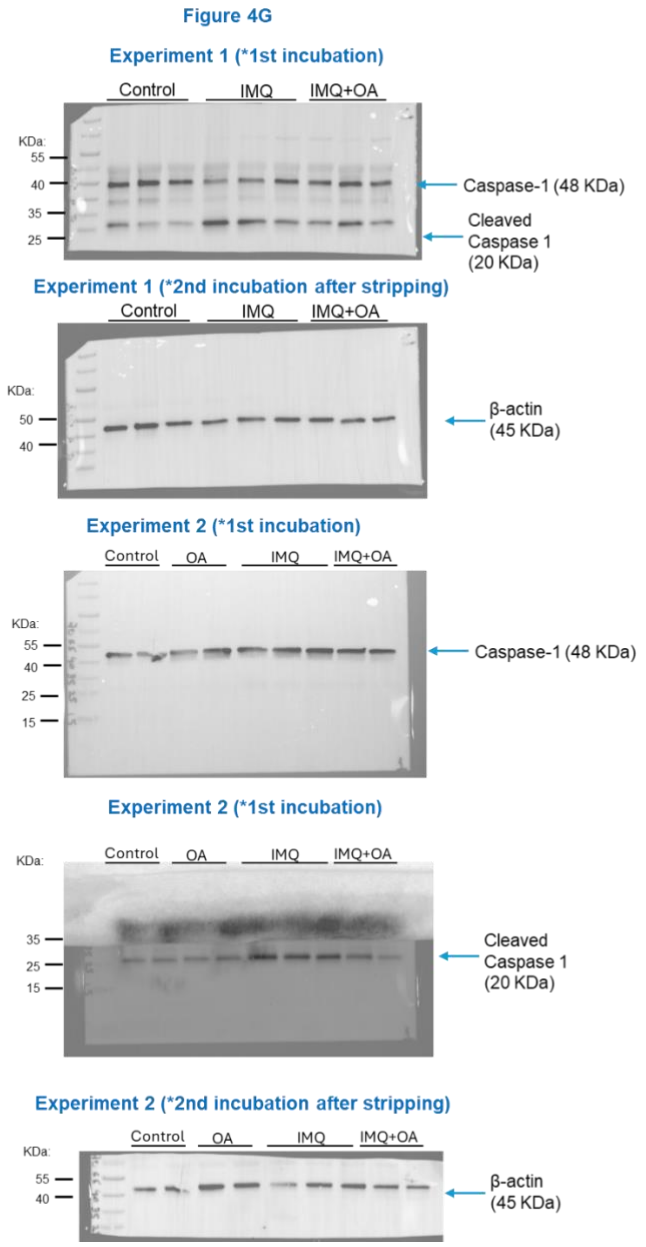


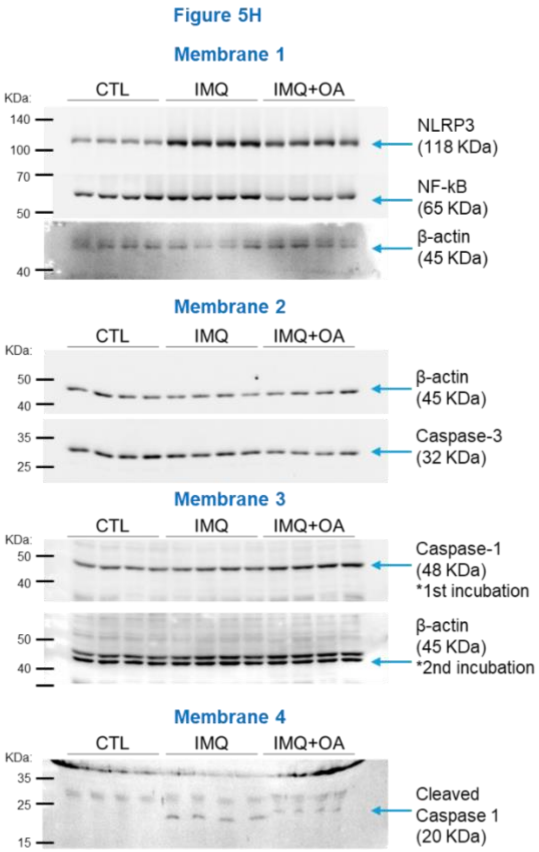


Uncropped, full-length images of the Western blot membranes presented in the main Figures 4G and 5H.

In the images of Figure 4G, the “OA” group corresponds to skin samples collected from mice supplemented with oleic acid without the treatment with IMQ; however, these samples were not included in the comparative analyses. The samples shown in Figure 4G were analysed using two separate membranes in two independent experiments.
